# Bilateral JNK activation is a hallmark of interface contractility and promotes elimination of aberrant cells

**DOI:** 10.1101/2022.06.20.496791

**Authors:** Deepti Prasad, Katharina Illek, Friedericke Fischer, Kathrin Holstein, Anne-Kathrin Classen

## Abstract

Tissue-intrinsic defence mechanisms eliminate aberrant cells from epithelia and thereby maintain the health of developing tissues or adult organisms. ‘Interface contractility’ comprises one such distinct mechanism that specifically guards against aberrant cells, which undergo inappropriate cell fate and differentiation programs. The cellular mechanisms which facilitate detection and elimination of these aberrant cells are currently unknown. We find that in *Drosophila* imaginal discs, interface contractility is associated with bilateral JNK activation at the clonal interface of wild type and aberrant cells. Bilateral JNK activation is unique to interface contractility and is not observed in other tissue-intrinsic defence mechanisms, such as cell-cell competition. We find that JNK is activated cell-autonomously by either of the contacting cell types and drives apoptotic elimination of cells at clonal interfaces. Ultimately, JNK interface signalling provides a collective tissue-level mechanism, which ensures elimination of aberrant, misspecified cells that cannot be identified by cell fitness criteria, as in cell-cell competition. Importantly, oncogenic Ras activates interface contractility but evades apoptotic elimination by bilateral JNK activation. Combined, our work establishes bilateral JNK interface signalling and interface apoptosis as a new hallmark of interface contractility, and it highlights how oncogenic mutations evade tumour suppressor function encoded by this tissue-intrinsic surveillance system.

## Introduction

Genetically altered cells appear in epithelial tissues at constant rate, either as a result of developmental errors [1] or mutagenesis throughout adult life [2]. Surveillance and removal of genetically altered cell is required to maintain tissue and organismal health. In addition to immune cell dependent processes [3–5], tissue-intrinsic mechanisms, such as cell-cell competition, epithelial defence against cancer (EDAC) or interface contractility, have been identified to remove aberrant cells from tissues [6–11]. In classical cell-cell competition scenarios, the comparison of cell fitness between neighbouring cells is the driving force of cell elimination, with less fit ‘loser’ cells being eliminated by fit ‘winner’ cells [9, 10, 12–14]. Mutations that interfere with house-keeping functions, such as proteostasis, cellular metabolism or genome maintenance, have emerged as drivers of cell-cell competition [13, 15–21].

In contrast, mutations may also interfere with signalling pathways or transcriptional networks that set up specific cell fate and differentiation programs [22–24]. These mutations may change a cell’s fate trajectory but importantly, may not disrupt cellular fitness *per se*. In the absence of fitness information linked to a genotype, these aberrant misspecified cells create a distinct challenge to tissue health and, consequently, must activate a distinct programme for their detection and elimination from tissue. Such programme must be able to deal with egalitarian cell fate information at cell contact sites, as neither normal nor aberrant cell fate genotypes carry inherent information about which of the two neighbouring cells should survive. A decision about which cell type is aberrant can thus only be determined though a potentially meta-democratic process where a read-out for ‘aberrant’ is generated by a collective decision of many cells, rather than by comparison of fitness states between just two neighbouring cells, as in cell-cell competition.

We previously established a novel paradigm for tissue-intrinsic defence mechanisms against aberrant, misspecified cells in *Drosophila* imaginal discs that we termed ‘interface contractility’ [11]. Briefly, we demonstrated that two neighbouring cell types identify relative differences in cell fate and differentiation and react by recruiting actomyosin to shared contact interfaces. This drives cell segregation between two clonal cell populations via smoothening of the contractile interface and is accompanied by elevated apoptosis [11]. Interface contractility is a surprisingly universal response to relative differences in cell fate and differentiation states. Mosaic manipulation of the patterning pathways Dpp/TGF-β, Wg/WNT, Hh/Shh, JAK/STAT and Notch, or cell-fate-specifying transcription factors (Arm, APC, Iro-C, Omb, Yki, En/Inv, Ap, Ci) induces clone smoothening and apoptosis in imaginal discs [25–43]. Importantly, interface contractility is induced in a strict position-dependent manner according to the cell fate of the surrounding cells. For example, ci-expressing clones have normal corrugated shapes in anterior wing compartments, where ci-activity is high. However, ci-expressing clones undergo clone smoothening and die in posterior compartments, where ci-signalling is low [11]. Thus, interface contractility is exclusively driven by relative differences in cell fate programs between neighbouring cells where ‘aberrant’ is defined collectively in comparison to the program of the surrounding ‘normal’ cells. Indeed, we find that not only aberrant clones are eliminated from the tissue, but that wild type cells are also eliminated when surrounded by a majority of aberrant cells [11]. This distinguishes interface contractility from cell-cell competition, which responds to a clearly defined fitness gradient between two neighbouring cells, which ensures that always the aberrant loser cell dies, independent of spatial context. Interface contractility instead drives the elimination of a population that is in the minority with respect to the surrounding cells. As an extension of this principle, we described that elimination by interface contractility depends on clone size. Single aberrant cells and small aberrant clones are more efficiently eliminated. Moreover, small clones present with higher level of apoptosis than larger clones [11]. Yet, the molecular and cellular pathways, which drive elimination by interface contractility are currently not known.

Using *Drosophila* wing imaginal disc model, we describe here a surprisingly ingenious solution to remove aberrant cells by interface contractility. We find that interface contractility is associated with bilateral activation of pro-apoptotic JNK signalling interface-contacting cells of two differently fated cell populations. This egalitarian, cell-autonomous response of two contacting cells drives apoptosis at clonal interfaces. The topology of cell contacts, which depends on clone size, is consistent with a model whereby bilateral JNK activation underlies a collective decision about which of two genotypes represents a cell fate minority and thereby drives efficient elimination of single aberrant cells and small clones.

## Results

To better understand how aberrant cells die by interface contractility, we analysed cell elimination in mosaic wing imaginal discs (Fig. 1A-C) using two genotypic classes affecting cell fate: (1) cell fate regulators that are not normally expressed in the wing disc, and (2) those that are but in specific spatial pattern. The first group is represented by the transcription factors Fkh and Ey, which are master regulators of salivary gland [44, 45] and eye specification [46], respectively. These genes are not expressed in the wing discs (Fig.1E, Fig. S1A-D) and their ectopic expression causes interface contractility hallmarks, such as interfacial actomyosin enrichment and clone smoothening, at any position in the wing disc (Fig. 1D, F, Fig. S1C,E). The second group of cell fate programs is represented by Dpp-mediated (Tkv) (Fig. 1G,H), EGF-mediated (Egfr) (Fig. 1J,K, Fig. S1F,G), Wg-mediated (Arm) (Fig. S1I-K) and Hh-mediated (Ci) (Fig. S1M,N) patterning programs. Dpp, EGF and Wg-signalling are predominantly active in the central domain of the wing pouch, whereas Hh-signalling is active in the anterior compartment of the wing disc. Consequently, ectopic activation of these pathways causes interface contractility hallmarks, such as actomyosin enrichment and smoothening of clone boundaries, in more lateral pouch domains for expression of constitutively activate *tkv^CA^*, *Egfr^CA^* and *arm^S10^* constructs (Fig. 1 I,L; Fig. S1H,L), and in the posterior compartment for expression of *ci* (Fig. S1O). In agreement with the idea that the absolute difference between neighbouring cells induces interface contractility, we confirmed that conversely, loss of *tkv* function induces actomyosin enrichment and clone smoothening in domains where Dpp-signalling is usually high (Fig. S1P,Q). Similarly, previous studies demonstrate pattern-specific clone smoothening or even cyst formation for mosaic discs carrying LOF mutations in *vg* [27], *Iro-C* [42], *omb* [43], *ci* [47], *ap* [48], *en* [49] or Polycomb complexes [33]. Combined, these data highlight that aberrant cells, as identified by relative mispositioning with respect to the fate of surrounding cells, induce interface contractility hallmarks.

**Figure 1.**
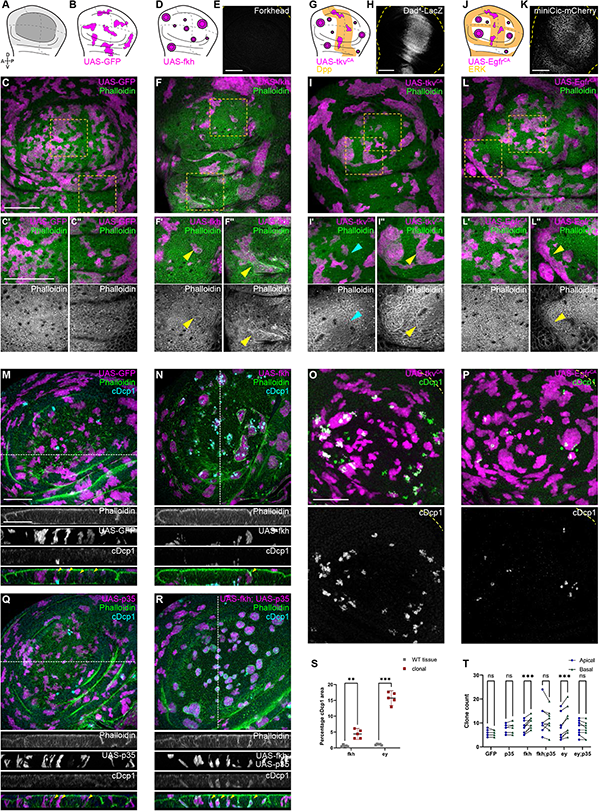
Apoptosis is essential to eliminate cells by interface contractility. **A:** Wing disc scheme highlighting the pouch (dark grey), hinge (light grey) and notum (white), as well as folds (continuous lines) and dorso-ventral or anterior-posterior compartment boundaries (dashed lines) (A). **B,D,G,J**: Wing discs schemes illustrating patterns of Fkh expression (D), Dpp signalling (G) or ERK activity (J) (orange represents endogenous expression or activity, white represents lack thereof). Clones (magenta) that do not induce interface contractility, because they only express GFP (B) or whose fate is like that of surrounding cells (magenta clones in orange domains in G, J), maintain irregular clone shapes. Clones whose fate is very different to surrounding cells, because of changes to cell fate and differentiation programs (magenta clones in white domains in D, G, J) experience interface smoothening and even cyst formation, as the apical surface buckles from interface contractility-induced compression. Clones express GFP (B), Fkh (D), Tkv^CA^ (G) and Egfr^CA^ (J). **E,H,K**: Wing discs stained to visualize endogenous patterns of Fkh expression (E), Dpp signalling (Dad^4^-LacZ, Dpp target gene) (H), and ERK signalling (*tub-miniCic-mCherry,* a ERK-activity reporter) (K). Yellow dashed lines indicate wing disc outlines. **C,F,I,L**: Wing disc carrying mosaic clones (magenta) expressing GFP (C), Fkh (F), Tkv^CA^ (I) and Egfr^CA^ (L) stained with phalloidin to visualize Actin (green or grey). Yellow frames mark regions in pouch centre (C’, F’, I’, L’), pouch periphery (I’’,L’’), and hinge (C’’,F’’). Cyan arrows point to accumulation of actin normally observed at the A-P compartment boundary (I’). Yellow arrows point to apical enrichment of actin at clone boundaries. (F’) focusses on the apical region of a clone that has undergone buckling. **M,N,Q,R**: Maximum-intensity projections of basal sections of wing discs carrying mosaic clones (magenta or grey) expressing GFP (M), Fkh (N), p35 (Q), and Fkh,p35 (R), stained with phalloidin to visualize Actin (green or grey) or for cDcp1 (cDcp1) to visualize apoptosis (cyan or grey). Dashed white lines indicate position of cross-sections. Yellow arrow heads in the cross-section overlays indicate viable clones still integrated in the epithelium. **O,P**: Maximum-intensity projections of basal sections of wing discs carrying mosaic clones (magenta) expressing Tkv^CA^ (O) or Egfr^CA^ (P) and stained for cDcp1 to visualize apoptosis (green or grey). Yellow dashed lines indicate wing disc outlines. Please refer to (G,J) for endogenous activation patterns of Dpp and ERK signalling in wing discs. **S:** Quantification of the percentage of apoptotic area in Fkh and Ey-expressing clones, as compared to the surrounding wildtype tissue. Graph displays mean± 95% CI. n=5 wing discs per genotype. **T:** Quantification of the number of clones detected apically (blue) or basally (green) in the wing disc pouch, where clones either express GFP; p35; Fkh; Fkh, p35; Ey or Ey, p35. n=9 wing discs per genotype. S, T: One-way ANOVA tests were performed to test for statistical significance, ns = not significant, **p≤0.01, ***p≤0.001. Scale bars = 50µm.

We previously reported a strong dependency of cell elimination on activation of apoptosis for ectopic expression of *fkh* [11]. When we analysed the distribution of apoptotic patterns in all genotypes, we observed that cDcp-1 positive events in *fkh-*, *ey-*, *tkv^CA^*-, *tkv-RNAi* and *Egfr^CA^*-expressing clones specifically occurred in positions, where also interface contractility hallmarks, such as clone smoothening and interphase actomyosin enrichment, occurred (Fig. 1M-P,S; Fig. S1R,S). Similarly, *ci-*expressing clones are specifically eliminated from posterior compartments [11]. Thus, elimination of mispositioned aberrant cells strongly correlates with interface contractility responses and suggests that cell elimination proceeds via apoptosis. To confirm that apoptosis was generally required to eliminate mispositioned cells from the tissue, we co-expressed p35, a strong inhibitor of apoptosis. Consequently, *fkh-* and *ey-*expressing clones survived and remained integrated in the epithelial layer (Fig. 1Q,R,T; Fig. S1T). We never observed basally delaminated cells, suggesting that mechanically driven live cell extrusion, as described for mammalian EDAC mechanisms [50, 51], is not a contributing process. As reported previously [11, 26, 27], inhibition of apoptosis did not interfere with clone smoothening nor with cyst formation, demonstrating that these morphological changes occur upstream or independent of apoptotic elimination. Combined, these results demonstrate that interface contractility defences activate apoptotic pathways to eliminate cells.

To understand how aberrant clones die by apoptosis, we analysed signalling through a potent tissue stress response pathway with important functions in cell death decision, namely JNK/AP-1 [52–54]. We found JNK to be consistently activated but, surprisingly, in a striking bilateral pattern at clonal interfaces. Specifically, in *fkh-* and *ey-*expressing clones, one cell row on each side of the interface displayed activation of the JNK-reporter *TRE-RFP* (Fig. 2A-E; Fig. S2.1A,B; Fig. S2.2A,B) [55]. The reporter *puc-LacZ*, while significantly less sensitive, independently confirmed that bilateral interface signals were JNK-specific (Fig. S2.3). This indicates that both wild type and aberrant cells activate JNK signalling when in contact with each other. Consistent with the idea that interface contractility strictly responds to relative fate differences between neighbouring cell, we also observed bilateral JNK activity when *fkh-* or *ey-*expressing cells represent the majority population in the tissue (Fig. 2F-J; Fig. S2.1 C,D; Fig. S2.2 C,D). Importantly, bilateral JNK interface responses could also be observed in discs carrying *tkv^CA^*, *Egfr^CA^*, *arm^S12^* and *ci*-expressing clones (Fig. 2K,L; Fig. S2.1 E-I; Fig. S2.2E-J). Here, however, JNK interface signalling strictly correlated with position of clones within the respective Dpp-, EGF-, Wg- and Hh-patterning field, confirming that JNK interface signalling responds to relative cell fate differences between neighbouring cells. Combined, these observations establish JNK interface signalling as novel hallmark of interface contractility defences and highlights the activation of a central stress signalling pathway upon direct surface contact between distinct cell fates.

**Figure 2.**
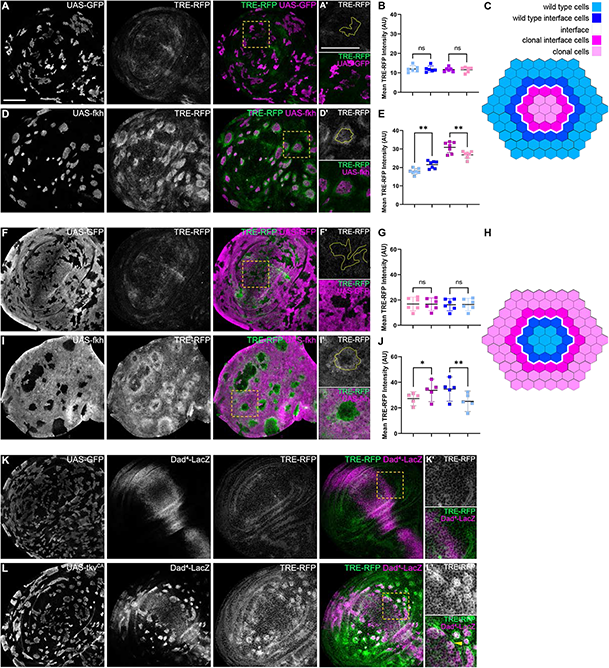
Interface contractility activates bilateral JNK interface signalling. **A,D,F,I**: Wing discs carrying mosaic clones (magenta or grey) expressing GFP (A,F) or Fkh (D,H), as well as the JNK reporter TRE-RFP (green or grey). Yellow frames mark regions shown in (A’, D’, F’, H’,). Continuous yellow line in (‘) panels marks clone boundaries. **C,H**: Schemes depicting specific zones in and around clones that were quantified. **B,E,G,J**: Quantifications of TRE-RFP intensities in specific zones of clones expressing GFP (B) or Fkh (E), and GFP-negative wild type clones amidst GFP-expressing wild type cells (G), and GFP-negative wild type clones amidst Fkh-expressing cells (I). Graphs display mean± 95% CI. One-way ANOVA tests were performed to test for statistical significance, ns = not significant, *p≤0.05, **p≤0.01. n=6 wing discs (B,E,G) and n=5 wing discs (I). **K,L**: Wing disc carrying mosaic clones (grey) expressing GFP (J) or Tkv^CA^ (K) and the Dpp reporter Dad-LacZ (grey or magenta) and the JNK reporter TRE-RFP (grey or green). Yellow frames mark regions shown in (J’, K’). Dad-LacZ is induced by Tkv^CA^-expression, revealing where Tkv^CA^ clones are like their surroundings, and where they are not. Yellow arrows highlight TRE-RFP in wildtype cells at the interface. Scale bars = 50µm.

We wanted to first understand if bilateral JNK activation is a cell-autonomous response induced by apposition of different cell fates, or if it is based on non-autonomous signalling between the two cell types. We thus reduce JNK signalling within one cell type by expressing a dominant-negative *bsk* (*bsk^DN^*) construct in cells that also express *ey* or *tkv^CA^* (Fig. 3 A-D; Fig. S3 A,B). *bsk^DN^* is a potent suppressor of JNK signalling and completely abolished even strong activity of JNK in wild type and mispositioned clones, as judged by complete lack of intra-clonal *TRE-RFP* reporter activity (Fig. 3B,D; Fig. S3B) [56]. Strikingly, eliminating JNK signalling on the *ey-*expressing side of a clonal interface did not prevent JNK activation in neighbouring wild type cells, irrespective of whether these wild type cells enclosed *ey-*expressing cells, or were the ones being enclosed by *ey-*expressing cells (Fig. 3C-H). Similarly, expression of *bsk^DN^* did not prevent the Dpp pattern-specific activation of *TRE-RFP* in wild type cell at the interface with *tkv^CA^*-expressing clones (Fig. S3A-D). These experiments demonstrate that JNK interface signalling is activated cell-autonomously at the interface upon apposition of different cell fates. This is in agreement with a model where interface contractility responds to relative differences in cell fate, rather than vectorial and quantitative information such as lower or higher fitness.

**Figure 3.**
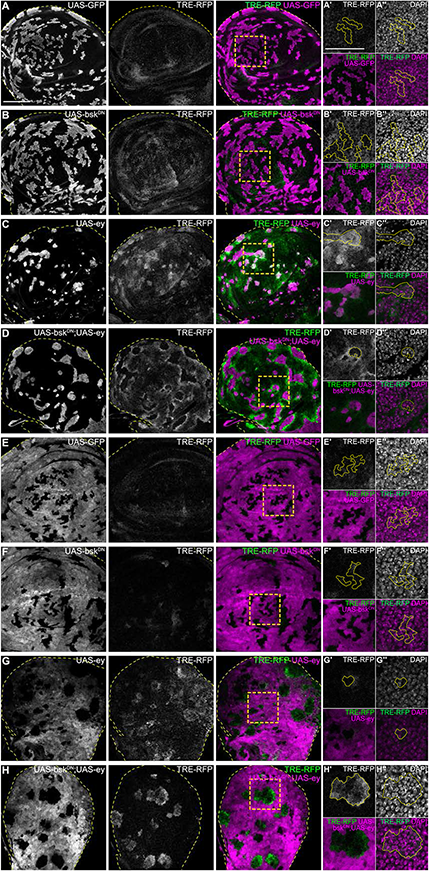
JNK activation is a cell-autonomous response to apposition of different fates. **A-H**: Wing discs carrying mosaic clones (magenta or grey) expressing GFP only (A, E); Bsk^DN^ (B, F), Ey (C, G), or Bsk^DN^, Ey (D, H) and expressing the JNK reporter TRE-RFP (grey or green). Discs were stained with DAPI to visualize individual nuclei (grey) (A’’-H’’)). Yellow dashed lines in (A-H) demarcate the wing disc outline. Yellow frames mark regions shown in (‘) and (’’) panels. Continuous yellow line in (‘ and’’) panels marks clone boundaries. Scale bar = 50µm.

Importantly, these experiments also confirmed that all wing discs domains have the competence to activate JNK interface signalling. Normally, mispositioned clones in the pouch were smaller, apoptotic and with high clone-intrinsic levels of JNK activity (Fig. 1N, Fig.2D, Fig. S2.2A). This partially confounded the resolution of bilateral JNK interface pattern. However, larger clones in the pouch area obtained by expression of *bsk^DN^* displayed JNK activation in surrounding wild type cells (Fig.3D, Fig. S3B). Thus, while the resolution and strength of JNK activation may be affected by tissue-specific dynamics, all areas of the wing disc can respond with JNK activation at the interface to the presence of mispositioned cells.

To understand if JNK interface signalling was a specific hallmark of interface contractility, we analysed JNK-reporter activity in mosaic tissues subject to classical cell-cell competition. We specifically analysed three well-established cell-cell competition genotypes: (1) clones mutant for *RpS3*, representing a loser genotype [57], (2) clones ectopically expressing *Dmyc* [58, 59] and (3) clones with reduced levels of the Hippo/Yorkie pathway component *wts* [60], both representative super-competitors. Importantly, none of these three genotypes displayed JNK interface signalling (Fig. 4, Fig. S4.1). Thus, while JNK activity may be observed cell-autonomously in loser genotypes due to disruption of cellular homeostasis [14], cell-cell competition does not induce JNK interface signalling. These results strongly demonstrate that JNK interface signalling is a specific hallmark of interface contractility and confirm that interface contractility is a tissue defence program that is distinct from canonical cell-cell competition.

**Figure 4.**
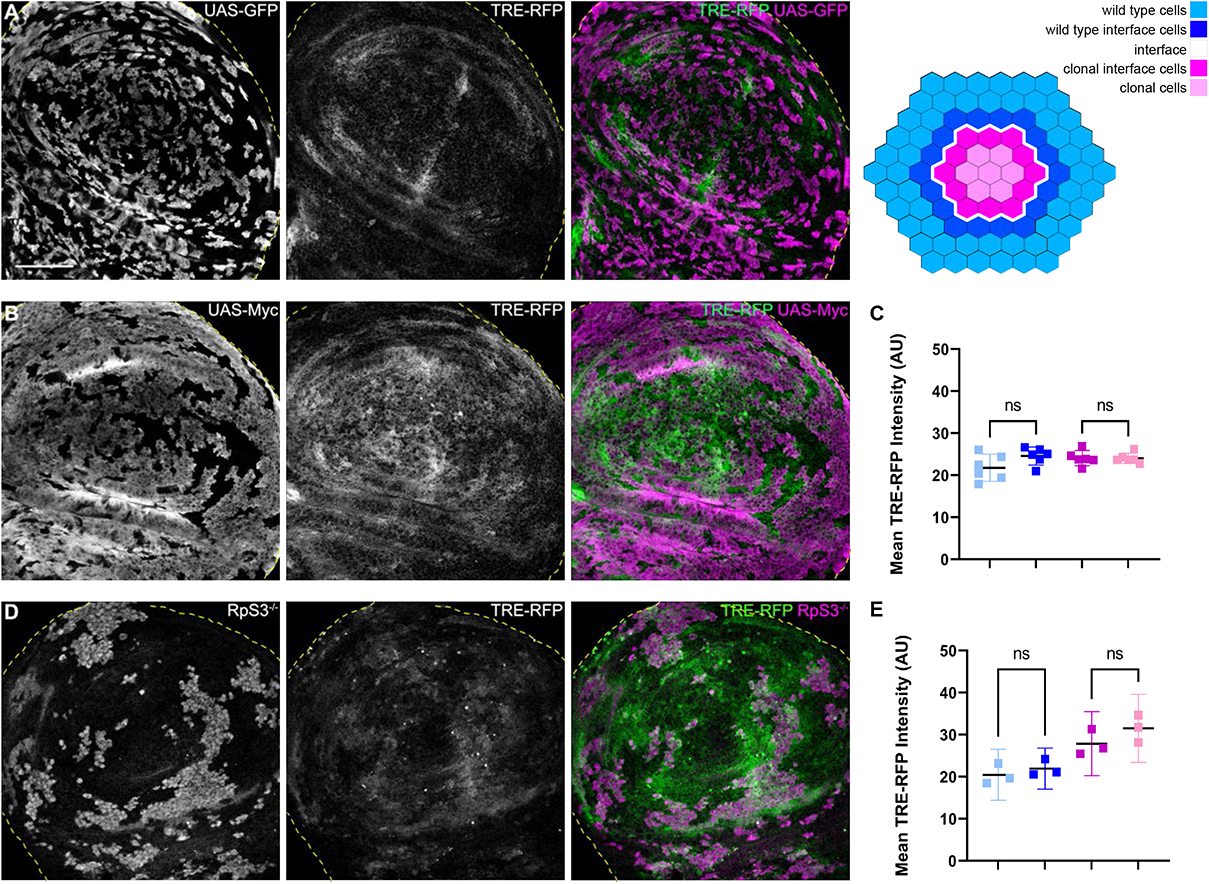
JNK interface signalling is unique to interface contractility and is not activated by cell competition. **A,B,D**: Wing disc carrying mosaic clones (grey or magenta) expressing GFP (A), Myc (B), or clones homozygous mutant for RpS3 (D). Discs express the JNK reporter TRE-RFP (grey or green). Yellow dashed lines demarcate the wing disc outline. **C,E**: Quantifications of TRE-RFP intensities in clones expressing Myc (C), or in clones homozygous mutant for *RpS3* (E). Graphs display mean± 95% CI. One-way ANOVA tests were performed to test for statistical significance, ns = not significant. n=6 wing discs (C), n=3 wing discs (E). Note that the background TRE-RFP in RpS3^-/-^ mosaic discs was measured in a 4 µm zone outside of wild type interface cells, as TRE-RFP was broadly activated in these discs. Scale bars = 50µm.

Another hallmark that distinguishes interface contractility from cell-cell competition is enrichment of actomyosin at clonal interfaces, a feature not observed in classical cell-cell competition. The co-occurrence of JNK signalling and actomyosin enrichment at the clonal interface, and the fact that JNK can be an upstream regulator of the actin dynamics [61–63], led us to ask if JNK interface signalling recruits actomyosin. To test this, we first reduced JNK activity within clones by co-expression of a dominant-negative *bsk* (*bsk^DN^*) construct. However, the interfaces of ey, *bsk^DN^* or *tkv^CA^*, *bsk^DN^*-expressing clones remained smooth, and cyst formation as well as actin enrichment could still be observed (Fig. S4.2A-D). To test the contribution of JNK interface signalling at both side of the clonal interface, we expressed *bsk^DN^* in the entire posterior compartment and induced *tkv^CA^*-expressing clones. However, no changes to actin enrichment or smoothness of interfaces could be observed in *tkv^CA^* clones (Fig. S4.2E,F). Combined, these observations establish that JNK interface signalling acts in parallel or downstream of elevated actomyosin contractility at clonal interfaces and thus serves another independent function in interface contractility defences.

Because JNK signalling can act as a promoter of apoptosis, we asked if JNK interface signalling serves to promote the elimination of cells, specifically at the interface. To test this idea, we first analysed the spatial distribution of apoptotic events in different mosaic genotypes (*fkh, ey* and *tkv^CA^*). To distinguish apoptotic events at the interface from apoptotic events in the clone interior, we specifically analysed large clones with separable interface, interior and exterior zones (Fig. S5A). This analysis revealed that many, but not all apoptotic events, occurred at clonal interfaces in *fkh-, ey-* or *tkv^CA^-*expressing clones (Fig. 5A-F; Fig. S5B,C). Importantly, apoptotic events were also elevated in wild type interface cells, indicating that JNK interface signalling also affects the contacting wild type cells.

**Figure 5.**
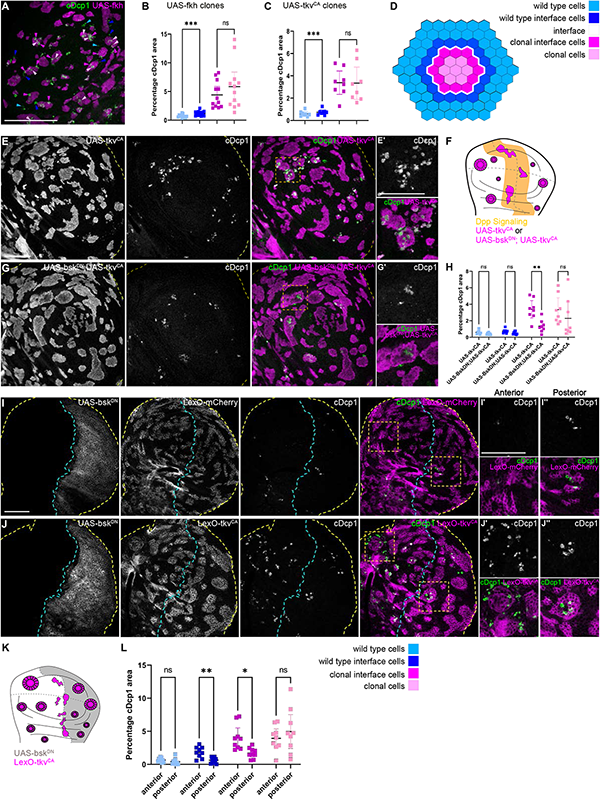
Cell elimination at clonal interfaces is mediated by JNK. **A**: Maximum-intensity projection of basal sections from wing imaginal discs carrying clones (magenta) expressing Fkh and stained for cDcp1 to visualize apoptosis (green). Coloured arrows show apoptosis in different zones that were used for quantification: light blue (wild type cells), dark blue (wildtype interface cells), magenta (clonal cells at interface), light pink (clonal cells). **B,C**: Quantifications of cDcp1 area fractions in selected zones around clones expressing UAS-Fkh (B) or UAS-Tkv^CA^ (C). Only large clones with distinct interior zones were quantified for clone interior and clonal interface zones. Graphs display mean± 95% CI. One-way ANOVA tests were performed to test for statistical significance, ns = not significant, *p p≤0.05, ** p≤0.01, *** p≤0.001. n=12 wing discs (B) and n=8 wing discs (C). **D**: Scheme depicting specific zones in and around clones that were quantified. **E,G**: Maximum-intensity projections of basal sections of wing imaginal discs carrying clones (grey or magenta) expressing Tkv^CA^ (A), or Bsk^DN^,Tkv^CA^ (B) stained for cDcp1 to visualize apoptosis (grey or green). Yellow dashed lines demarcate the wing disc outline. Yellow frames mark regions shown in (A’, B’). **F:** Scheme illustrates endogenous patterns of Dpp signalling (orange) and expected smoothening of Tkv^CA^ and Tkv^CA^, Bsk^DN^ expressing clones (magenta) in areas where Dpp signalling is not normally active (white). **H:** Quantifications of cDcp1 area fractions in selected zones around clones expressing UAS-Tkv^CA^ or UAS-Bsk^DN^,UAS-Tkv^CA^. Only large clones with distinct interior zones were quantified for clone interior and clonal interface zones. Graphs display mean± 95% CI. Two-way ANOVA tests were performed to test for statistical significance, ns = not significant, * p≤0.05, ** p≤0.01. n=8 wing discs per genotype. **I,J**: Maximum intensity projection of basal section of wing discs, where the posterior compartment expresses Bsk^DN^ (grey) under the control of en-GAL4, and where clones (grey or magenta) express LexO-mCherry (E) or LexO-Tkv^CA^ (F). Discs were stained for cDcp1 to visualize apoptosis (grey or green). Yellow dashed lines demarcate the wing disc outline. Yellow frames mark regions selected in anterior (E’, F’) and posterior (E’’, F’’) compartments. Dashed cyan line highlights the anterior-posterior compartment boundary. **K:** Scheme illustrates expression of Bsk^DN^ in the posterior compartment and the response of Tkv^CA^ expressing clones (magenta). **L**: Quantifications of cDcp1 area fractions in selected zones around clones expressing LexO-Tkv^CA^ either in the anterior control compartment or in the posterior compartment expressing Bsk^DN^. Only large clones with distinct interior zones were quantified for clone interior and clonal interface zones. Graphs display mean± 95% CI. Two-way ANOVA tests were performed to test for statistical significance, ns = not significant, * p≤0.05, ** p≤0.01. n=9 wing discs. Scale bar = 50µm

To test if JNK interface signalling was indeed mediating cell elimination of aberrant cells, we asked how inhibition of JNK may change the spatial distribution of apoptotic events. We utilized two genetic strategies: we first inhibited JNK by expression of *bsk^DN^* within *tkv^CA^*- expressing clones (Fig. 5F-H) and secondly, by expression in the posterior compartment of a disc harbouring *tkv^CA^* clones (Fig. 5I-L; Fig. S5D). Strikingly, inhibition of JNK strongly reduced apoptosis at intra-clonal interfaces. Importantly, we also observed a reduction in apoptosis in interface wild type cells, demonstrating that wild type cell survival is also affected by JNK interface signalling. The observation that both wild type and aberrant cell survival is regulated by JNK interface signalling complements the idea that interface contractility defences act upon local differences in cellular programs rather than on a specific cell identity *per se*. Combined, these results clearly establish that JNK interface signalling strongly contributes to the elimination of cells at clonal interfaces and is thus a functional hallmark of the interface contractility program.

Intriguingly, JNK interface signalling cannot account for all apoptotic events that we observed in aberrant clones. A second population of apoptotic cells, which could not be suppressed by expression of *bsk^DN^*, located specifically to interior zones of cystic clones. These interior zones have lower JNK activity (Fig. 2E, J; Fig. S2.2 B,D) but their apical surface often buckles to form cysts [11]. Theoretically, cysts arise when the contractile interface compresses small and intermediate-sized clones, thereby driving apical surface buckling and cyst formation [11]. When we analysed apoptotic patterns, we observed a strong spatial correlation between apical buckling and the apical presence of a subpopulation of apoptotic cells (Fig. S5E-G). This raises the possibility that cell death in clone interiors may be driven by surface buckling dynamics and likely represents the source of JNK-independent apoptotic cell death (Fig S5.1). While this needs to be investigated further, we conclude that two spatially distinct and independent mechanisms act to promote elimination of aberrant cells.

Mutations in Wnt, TGFD or cell fate specification pathways are widely reported to promote cancer [22–24]. We found that mutations in these pathways consistently elicit interface contractility and ultimately, apoptosis of affected cells. Thus, interface contractility would play an important tumour suppressive role by facilitating elimination of cells with aberrant fate. One exception was revealed by our analysis of *Ras^V12^*-expressing clones, representing an incredibly potent oncogenic mutations in cancer patients [64]. *Ras^V12^* cells activate ERK signalling at high levels (Fig. S6.1A,B,F) and exhibit ERK pattern-specific actomyosin enrichment and JNK activation at the interface with surrounding wild type cells (Fig. 6A,B). Thus, *Ras^V12^*-expressing clones activated hallmarks of interface contractility and were recognized as a distinct cell fate. However, *Ras^V12^*-expressing clones completely failed to induce apoptosis and were not eliminated from imaginal discs, suggesting that *Ras^V12^* potently suppresses JNK interface and buckling-associated apoptosis (Fig. 6C,D,G; Fig. S6.2A). Moreover, *Ras^V12^* could dominantly suppress apoptosis in *fkh*- and *ey-*expressing clones, likely via dominantly elevating ERK activity (Fig. 6E-G; Fig. S6.1C-E,G,H; Fig. 6.2B,C). *Ras^V12^* is the only defined genotype for which we have observed complete evasion of apoptotic elimination by the interface contractility program, highlighting the central role of oncogenic Ras as an initiating and cooperating factor in tumour formation. In conclusion, while misspecification of cell fate induces interface contractility, oncogenic mutations may escape this tissue-intrinsic defence by promoting apoptosis resistance, thus increasing the likelihood of clone persistence and thus tumour growth.

**Figure 6.**
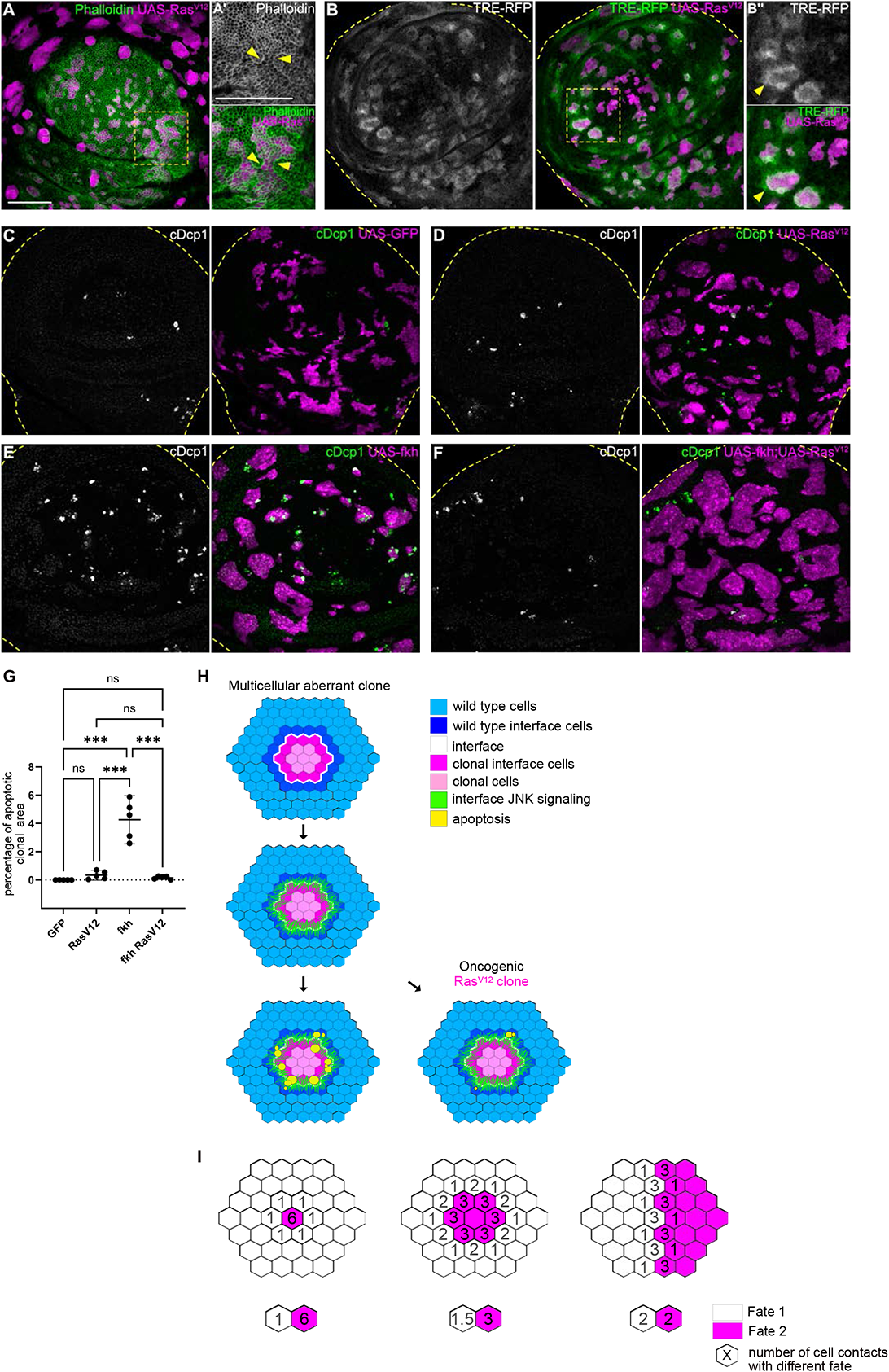
Cell elimination by interface contractility is suppressed by oncogenic Ras^V12^. **A**: Wing disc carrying mosaic clones (magenta) expressing Ras^V12^ were stained with phalloidin to visualize Actin (grey or green). Yellow frames mark regions shown in (A’). Yellow arrows point to actin enrichment at clone boundaries. **B**: Wing disc carrying mosaic clones (magenta) expressing Ras^V12^ and TRE-RFP (grey or green). Yellow dashed lines demarcate the wing disc outline. Yellow frames mark regions shown in (B’). Yellow arrows highlight TRE-RFP activation. **C,D,E,F**: Maximum intensity projection of basal sections of wing discs carrying clones (magenta) expressing GFP (C) Ras^V12^ (D), Fkh (E), or Fkh, Ras^V12^ (F) were stained for cDcp1 to visualize apoptosis (grey or green). **G**: Quantifications of the percentage of apoptotic area in clones expressing GFP only, Ras^V12^, Fkh, or Fkh,Ras^V12^. Graphs display mean± 95% CI. One-way ANOVA tests were performed to test for statistical significance, ns = not significant, *** p≤0.001. n=5 wing discs per genotype. **H**: Model of bilateral JNK activation in cell elimination by interface contractility. Oncogenic mutations evade apoptosis. **I**: Model of how bilateral JNK activation drives cell elimination dependent on aberrant cell cluster size. Number of wild type-aberrant cell contacts determine the strength of pro-apoptotic JNK signalling. Scale bar = 50µm.

## Discussion

Here we demonstrate that cells of aberrant fate or differentiation state induce bilateral activation of JNK at clonal interfaces. We thus define bilateral JNK activation as an important new hallmark of interface contractility defences against aberrant cells. Indeed, activation of JNK has been previously observed when steep differences in cell fates occur [65, 66]. Bilateral JNK activation is consistent with a model where interface contractility monitors tissue health by assessing the degree of local differences and producing a bilateral response, rather than by linking apoptosis to a fixed loser genotype, as in cell-cell competition. Indeed, bilateral JNK activation promotes apoptosis in cells of either genotype that touch at the interface (Fig. 6H). So how could such a bilateral response create specificity for the elimination of aberrant cells? A specific mutation initially affects a single cell. All neighbouring wild type cells will induce bilateral JNK activation in the mutant cell, giving rise to high levels of JNK signalling in that cell. In contrast, any wild type cell will perceive bilateral JNK activation coming from just one mutant cell contact (Fig. 6I). As clone sizes increase, the number of interface junctions per aberrant cell decrease, and consequently, JNK activity as well. Yet, aberrant interface cells will still be in touch with more JNK-triggering contacts than surrounding wild type cells. In this model, bilateral JNK activation is particularly effective at eliminating single aberrant cells and small clones, which we indeed experimentally demonstrated previously [11]. Ultimately, the geometry of the interface contacts between two large cell populations equalizes and reduces the number of JNK-triggering cell contacts. As a consequence, larger clones are less like to die. The size-dependent geometry of bilateral JNK activation may also protect large fate compartments and their boundaries from high, potentially deleterious effects of JNK activation during development. Finally, while it may be surprising that evolution could accept that also interface wild type cells may occasionally be eliminated because of bilateral JNK activation, the advantage of such a general detector of cell fate mispositioning may outweigh the disadvantage of cell loss.

Strikingly, oncogenic *Ras^V12^* expression induces interface contractility hallmarks, such as interface actomyosin, buckling and cyst formation [11], as well as JNK interface signalling. However, *Ras^V12^* cells completely evade apoptosis, highlighting the potent oncogenic survival provided by *Ras^V12^* cooperativity. Importantly, expression of oncogenic Ras induces similar hallmark in mammalian MDCK cell culture or organoids, such as enrichment of actomyosin at cellular interfaces [67–69] and elimination efficiency that depends clone-size [70, 71]. Indeed, JNK interface like signalling has been reported for oncogenic Src-expressing cells in organoid cultures [72]. Importantly, oncogenic Ras is a potent suppressor of apoptosis in flies and mammalian tissues [73]. While this prevents elimination of these cells from imaginal discs, these cells are eliminated from mammalian tissue, however importantly: by a cell death independent mechanism that involves strong activation of interfacial actomyosin activity, driving mechanical squeezing out of the tissue layer [68-70, 74-78]. We thus suggest that interface contractility and EDAC are evolutionarily conserved expression of the same ancient tissue-intrinsic defence system that specifically acts against cell fate deregulation.

## Author Contributions

Conceptualization DP, KI, AKC

Validation DP, KI, AKC

Investigation DP, KI, FF, KH, AKC

Writing DP, KI, AKC

Visualization DP, KI, AKC

Supervision AKC

Funding Acquisition AKC

## Acknowledgements

We thank the reviewers for critical comments on the manuscript. We thank the LIC facility at the University of Freiburg for technical help with imaging. We thank David Bilder, Dirk Bohmann, Suzanne Eaton, Iswar Hariharan, Martin Juenger, Romain Levayer, Giorgios Pyrowolakis and Helena Richardson for sharing reagents. We thank the Bloomington Drosophila Stock Centre (BDSC), the Vienna Drosophila Stock Collection (VDRC) and the Developmental Studies Hybridoma Bank (DSHB) for providing fly stocks and antibodies.

## Experimental Procedures

### Fly genetics

A list of strains, detailed genotypes and experimental conditions are provided in Supplemental Tables S1 and S2. Briefly, all crosses were kept on standard media. FLP/FRT and ‘GAL4/UAS flip-out’ and ‘LexA/LexO flip-out’ mosaic experiments utilized heat-shock-driven expression of a flipase. The respective crosses were allowed to lay eggs for 72 h at 25°C followed by a heat-shock at 37°C for 60 min (FLP/FRT) or 8-25 min (‘flip-out’). Larvae were dissected at wandering 3rd instar stage or as indicated (30 h, 54 h after heat-shock).

**Table S1.** Fly strains

| Genotype | Chrom. | Source |
| --- | --- | --- |
| *w<sup>118</sup>* | I | David Bilder |
| *hsflp<sup>122</sup>* | I | Iswar Hariharan |
| *act>y<sup>+</sup>>GAL4, UAS-GFP/TM6b* | III | David Bilder |
| *UAS-fkh-3xHA* | I | Martin Juenger |
| *hsflp<sup>122</sup>; Dad<sup>4</sup>-LacZ/CyO* | I,II,III | Giorgios Pyrowolakis |
| *UAS-tkv<sup>CA</sup>* | I | BDSC 36537 |
| *tub-miniCic-mCherry (or miniCic-mCherry)* | II | Romain Levayer |
| *tub-miniCic-mScarlett (or miniCic-mScarlett)* | II | Romain Levayer |
| *UAS-Egfr<sup>CA</sup>/TM6c* | III | BDSC 59843 |
| *UAS-p35* | II | David Bilder |
| *UAS-p35* | III | David Bilder |
| *UAS-ey* | II | BDSC 6294 |
| *UAS-arm<sup>S10</sup>* | I | Suzanne Eaton |
| *UAS-ci-HA* | III | BDSC 32570 |
| *UAS-tkv RNAi (TRiP.HMS02185)* | II | BDSC 40937 |
| *TRE-RFP* | II | Dirk Bohmann |
| *puc<sup>A251.1F3</sup>>LacZ/TM6c* | III | BDSC 11173 |
| *UAS-bsk<sup>DN</sup>* | I | BDSC 6409 |
| *UAS-myc-HA* | III | BDSC 64759 |
| *FRT82B ubi-GFP, RpS3<sup>Plac92</sup>/TM6c* | III | BDSC 5627 |
| *UAS-wts RNAi (101055/KK)* | II | VDRC 111002 |
| *en-GAL4, UAS-GFP* | II | David Bilder |
| *brk*-GAL4; *UAS*-CD8-GFP, *LexO*-mCherry-CAAX/SM5-TM6b; *LexO*-tkv<sup>CA</sup> /SM5-TM6b | I,II,III | Giorgios Pyrowolakis |
| *tub*-GAL80<sup>ts</sup> | III | BDSC 7018 |
| *UAS*-Ras<sup>V12</sup> | II | Helena Richardson |
| *UAS*-Ras<sup>V12</sup> | III | BDSC 4847 |

**Table S2.** Detailed genotypes

| Figure | Genotype | Heatshock | Analysis after: |
| --- | --- | --- | --- |
| 1 C,E,M | *hsflp*<sup>122/+</sup>; +/+;<br>*act>y<sup>+</sup>*>GAL4, *UAS*-GFP/+ | 9.5 min,<br>8.5 min | 30 h 25 °C |
| 1 F,N | *UAS-fkh-3xHA/hsflp*<sup>122</sup>; +/+;<br>*act>y<sup>+</sup>*>GAL4, *UAS*-GFP/+ | 9.5 min,<br>8.5 min | 30 h 25 °C |
| 1 H | *hsflp*<sup>122/+</sup>; *TRE-RFP/Dad*<sup>4</sup>-LacZ;<br>*act>y<sup>+</sup>*>GAL4, *UAS*-GFP/+ | 9.5 min | 30 h 25 °C |
| 1 I,O | *hsflp*<sup>122/+</sup>; +/+;<br>*act>y<sup>+</sup>*>GAL4, *UAS*-GFP/*UAS*-tkv<sup>CA</sup> | 9.5 min | 30 h 25 °C |
| 1 K | *hsflp*<sup>122/+</sup>; *tub-miniCic-mCherry*/+;<br>*act>y<sup>+</sup>*>GAL4, *UAS*-GFP/+ | 9.5 min | 30 h 25 °C |
| 1 L,P | *hsflp*<sup>122/+</sup>; +/+;<br>*act>y<sup>+</sup>*>GAL4, *UAS*-GFP/*UAS*-Egfr<sup>CA</sup> | 9.5 min | 30 h 25 °C |
| 1 Q | *hsflp*<sup>122/+</sup>; +/+;<br>*act>y<sup>+</sup>*>GAL4, *UAS*-GFP/*UAS*-p35 | 8.5 min | 30 h 25 °C |
| 1 R | *UAS-fkh-3xHA/hsflp*<sup>122</sup>; *UAS*-p35/+;<br>*act>y<sup>+</sup>*>GAL4, *UAS*-GFP/+ | 8.5 min | 30 h 25 °C |
| S1 A | *UAS-fkh-3xHA/hsflp*<sup>122</sup>; +/+;<br>*act>y<sup>+</sup>*>GAL4, *UAS*-GFP/+ | 9.5 min | 30 h 25 °C |
| S1 B,E,S | *hsflp*<sup>122/+</sup>; *UAS*-ey/+;<br>*act>y<sup>+</sup>*> GAL4, *UAS*-GFP/+ | 9.5 min,<br>8.5 min | 30 h 25 °C |
| S1D,J,K,N | *hsflp*<sup>122/+</sup>; +/+;<br>*act>y<sup>+</sup>*>GAL4, *UAS*-GFP/ + | 9.5 min | 30 h 25 °C |
| S1G | *hsflp*<sup>122/+</sup>; *tub-miniCic-mCherry*/+;<br>*act>y<sup>+</sup>*>GAL4, *UAS*-GFP/+ | 9.5 min | 30 h 25 °C |
| S1H | *hsflp*<sup>122</sup>/+; *tub-miniCic-mCherry*/+;<br>*act>y<sup>+</sup>>GAL4, UAS-GFP/UAS-Egfr<sup>CA</sup>* | 9.5 min | 30 h 25 °C |
| S1L | *UAS-arm<sup>S10</sup>/hsflp*<sup>122</sup>; +/+;<br>*act>y<sup>+</sup>>GAL4, UAS-GFP/+* | 9.5 min | 30 h 25 °C |
| S1O | *hsflp*<sup>122</sup>/+; +/+;<br>*act>y<sup>+</sup>>GAL4, UAS-GFP/UAS-ci-HA* | 9.5 min | 54 h 18 °C |
| S1Q,R | *hsflp*<sup>122</sup>/+; *UAS-tkv RNAi*/+;<br>*act>y<sup>+</sup>>GAL4, UAS-GFP/+* | 9.5 min | 30 h 25 °C |
| S1T | *hsflp*<sup>122</sup>/+; *UAS-ey*/+;<br>*act>y<sup>+</sup>>GAL4, UAS-GFP/UAS-p35* | 8.5 min | 30 h 25 °C |
| 2 A,F | *hsflp*<sup>122</sup>/+; *TRE-RFP*/+;<br>*act>y<sup>+</sup>>GAL4, UAS-GFP/+* | 8.5 min,<br>25 min | 30 h 25 °C |
| 2 D,I | *UAS-fkh-3xHA/hsflp*<sup>122</sup>; *TRE-RFP*/+;<br>*act >y<sup>+</sup>>GAL4, UAS-GFP/+* | 8.5 min,<br>25 min | 30 h 25 °C |
| 2 K | *hsflp*<sup>122</sup>/+; *TRE-RFP/Dad<sup>4</sup>-LacZ*;<br>*act>y<sup>+</sup>>GAL4, UAS-GFP/+* | 9.5 min | 30 h 25 °C |
| 2 L | *hsflp*<sup>122</sup>/+; *TRE-RFP/Dad<sup>4</sup>-LacZ*;<br>*act>y<sup>+</sup>>GAL4, UAS-GFP/ UAS-tkv<sup>CA</sup>* | 9.5 min | 30 h 25 °C |
| S2.1 A,C | *hsflp*<sup>122</sup>/+; *TRE-RFP*/+;<br>*act>y<sup>+</sup>>GAL4, UAS-GFP/+* | 8.5 min,<br>25 min | 30 h 25 °C |
| S2.1 B,D | *UAS-fkh-3xHA/hsflp*<sup>122</sup>; *TRE-RFP*/+;<br>*act >y<sup>+</sup>>GAL4, UAS-GFP/+* | 8.5 min,<br>25 min | 30 h 25 °C |
| S2.1 E | *hsflp*<sup>122</sup>/+; *TRE-RFP/Dad<sup>4</sup>-LacZ*;<br>*act>y<sup>+</sup>>GAL4, UAS-GFP/+* | 9.5 min | 30 h 25 °C |
| S 2.1 F | *hsflp*<sup>122</sup>/+; *TRE-RFP/Dad<sup>4</sup>-LacZ*;<br>*act>y<sup>+</sup>>GAL4, UAS-GFP/ UAS-tkv<sup>CA</sup>* | 9.5 min | 30 h 25 °C |
| S 2.2 A,C | *hsflp*<sup>122</sup>/+; *TRE-RFP/UAS-ey*;<br>*act>y<sup>+</sup>>GAL4, UAS-GFP/+* | 8.5 min,<br>25 min | 30 h 25 °C |
| S2.2 E,G,I | *hsflp*<sup>122</sup>/+; *TRE-RFP*/+;<br>*act>y<sup>+</sup>>GAL4, UAS-GFP/+* | 9.5 min | 30 h 25 °C |
| S2.2 F | *hsflp*<sup>122</sup>/+; *TRE-RFP*/+;<br>*act>y<sup>+</sup>>GAL4, UAS-GFP/UAS-Egfr<sup>CA</sup>* | 9.5 min | 30 h 25 °C |
| S2.2 H | *UAS-arm<sup>S10</sup>/hsflp*<sup>122</sup>; *TRE-RFP*/+;<br>*act>y<sup>+</sup>>GAL4, UAS-GFP/+* | 9.5 min | 30 h 25 °C |
| S2.2 J | *hsflp*<sup>122</sup>/+; *TRE-RFP*/+;<br>*act>y<sup>+</sup>>GAL4, UAS-GFP/UAS-ci-HA* | 9.5 min | 54 h 18 °C |
| S2.3 A | *hsflp*<sup>122/+</sup>; +/+; *puc*<sup>A251.1F3</sup>-*LacZ* /act>y<sup>+</sup>>GAL4, UAS-GFP/+ | 10 min | 30h 25 °C |
| S2.3 B | *UAS-fkh-3xHA/hsflp*<sup>122</sup>; +/+ ;<br>*puc*<sup>A251.1F3</sup>>*LacZ*/act>y<sup>+</sup>>GAL4, UAS-GFP | 10 min | 30 h 25 °C |
| S2.2 C | *hsflp*<sup>122/+</sup>; UAS-ey /+;<br>*puc*<sup>A251.1F3</sup>>*LacZ*/act>y<sup>+</sup>>GAL4, UAS-GFP/ + | 10 min | 30 h 25 °C |
| S2.3 D | *hsflp*<sup>122/+</sup>; + /+; *puc*<sup>A251.1F3</sup>-*LacZ*/act>y<sup>+</sup>>GAL4, UAS-GFP/UAS-Ras<sup>V12</sup> | 10 min | 30 h 25 °C |
| 3 A,E | *hsflp*<sup>122/+</sup>; TRE-RFP/UAS-ey;<br>act>y<sup>+</sup>>GAL4, UAS-GFP/+ | 9.5 min<br>25 min | 30 h 25 °C |
| 3 B,F | *UAS-bsk*<sup>DN</sup>/*hsflp*<sup>122</sup>; TRE-RFP/+;<br>act>y<sup>+</sup>>GAL4, UAS-GFP/+ | 9.5 min<br>25 min | 30 h 25 °C |
| 3 C,G | *hsflp*<sup>122/+</sup>; TRE-RFP/UAS-ey;<br>act>y<sup>+</sup>>GAL4, UAS-GFP/+ | 9.5 min<br>25 min | 30 h 25 °C |
| 3 D,H | *UAS-bsk*<sup>DN</sup>/*hsflp*<sup>122</sup>; TRE-RFP/UAS-ey;<br>act>y<sup>+</sup>>GAL4, UAS-GFP/+ | 9.5 min<br>25 min | 30 h 25 °C |
| S3A,C | *hsflp*<sup>122/+</sup>; TRE-RFP/+;<br>act>y<sup>+</sup>>GAL4, UAS-GFP/UAS-*tkv*<sup>CA</sup> | 9.5 min<br>25 min | 30 h 25 °C |
| S3B,D | *UAS-bsk*<sup>DN</sup>/*hsflp*<sup>122</sup>; TRE-RFP/+;<br>act>y<sup>+</sup>>GAL4, UAS-GFP/UAS-*tkv*<sup>CA</sup> | 9.5 min<br>25 min | 30 h 25 °C |
| 4 A | *hsflp*<sup>122/+</sup>; TRE-RFP/+;<br>act>y<sup>+</sup>>GAL4, UAS-GFP/+ | 9.5 min | 30 h 25 °C |
| 4 B | *hsflp*<sup>122/+</sup>; TRE-RFP/+;<br>act>y<sup>+</sup>>GAL4, UAS-GFP/UAS-*myc*-HA | 9.5 min | 30 h 25 °C |
| 4 C | *hsflp*<sup>122/+</sup>; TRE-RFP/+; FRT82B *ubi*-GFP,<br>*RpS3*<sup>Plac92</sup>/FRT82B | 1 h | 30 h 25 °C |
| S4.1 A | *hsflp*<sup>122/+</sup>; TRE-RFP/+;<br>act>y<sup>+</sup>>GAL4, UAS-GFP/+ | 9.5 min | 54 h 25 °C |
| S4.1 B | *hsflp*<sup>122/+</sup>; TRE-RFP/+;<br>act>y<sup>+</sup>>GAL4, UAS-GFP/UAS-*wt*s RNAi | 9.5 min | 54 h 25 °C |
| S4.2 A | *hsflp*<sup>122/+</sup>; TRE-RFP/UAS-ey;<br>act>y<sup>+</sup>>GAL4, UAS-GFP/+ | 9.5 min | 30 h 25 °C |
| S4.2 B | *UAS-bsk*<sup>DN</sup>/*hsflp*<sup>122</sup>; TRE-RFP/UAS-ey;<br>act>y<sup>+</sup>>GAL4, UAS-GFP/+ | 9.5 min | 30 h 25 °C |
| S4.2 C | *hsflp*<sup>122/+</sup>; TRE-RFP/+;<br>act>y<sup>+</sup>>GAL4, UAS-GFP/UAS-*tkv*<sup>CA</sup> | 9.5 min | 30 h 25 °C |
| S4.2 D | *UAS-bsk<sup>DN</sup>/hsflp<sup>122</sup>; TRE-RFP/+;*<br>*act>y<sup>+</sup>>GAL4, UAS-GFP/UAS-tkv<sup>CA</sup>* | 9.5 min | 30 h 25 °C |
| S4.2 E | *hsflp<sup>122</sup>/UAS-bsk<sup>DN</sup>; en-GAL4/+;*<br>*act>y<sup>+</sup>>UAS-GFP, LexO-mCherry /TM6c* | 9.5 min | 30 h 25 °C |
| S4.2 F | *hsflp<sup>122</sup>/UAS-bsk<sup>DN</sup>; en-GAL4/+;*<br>*act>y<sup>+</sup>>UAS-GFP, LexO-mCherry; LexO-*<br>*tkv<sup>CA</sup>/act>>VH2* | 9.5 min | 30 h 25 °C |
| 5 A | *UAS-fkh-3xHA/hsflp<sup>122</sup>; TRE-RFP/+;*<br>*act >y<sup>+</sup>>GAL4, UAS-GFP/+* | 8.5 min | 30 h 25 °C |
| 5 E | *hsflp<sup>122</sup>/+; TRE-RFP/+;*<br>*act>y<sup>+</sup>>GAL4, UAS-GFP/UAS-tkv<sup>CA</sup>* | 9.5 min | 30 h 25 °C |
| 5 F | *hsflp<sup>122</sup>/UAS-bsk<sup>DN</sup>; TRE-RFP/+;*<br>*act> y<sup>+</sup>>GAL4, UAS-GFP/UAS-tkv<sup>CA</sup>* | 9.5 min | 30 h 25 °C |
| 5 I | *hsflp<sup>122</sup>/UAS-bsk<sup>DN</sup>; en-GAL4/+;*<br>*act> y<sup>+</sup>>UAS-GFP, LexO-mCherry /TM6c* | 9.5 min | 30 h 25 °C |
| 5 J | *hsflp<sup>122</sup>/UAS-bsk<sup>DN</sup>; en-GAL4/+;*<br>*act> y<sup>+</sup>>UAS-GFP, LexO-mCherry; LexO-*<br>*Tkv<sup>CA</sup>/act>>VH2* | 9.5 min | 30 h 25 °C |
| S5 E | *hsflp<sup>122</sup>/UAS-fkh-3xHA; tub-GAL80<sup>ts-20</sup>/Sp or +;*<br>*act>y<sup>+</sup>>GAL4, UAS-GFP/Ly* | 11 min | 30 h 18 °C<br>18 h 30 °C |
| S5 F | *hsflp<sup>122</sup>/UAS-fkh-3xHA; tub-GAL80<sup>ts-20</sup>/Sp or +;*<br>*act>y<sup>+</sup>>GAL4, UAS-GFP/Ly* | 11 min | 30 h 30 °C |
| 6 A,D | *hsflp<sup>122</sup>/+; +/+;*<br>*act>y<sup>+</sup>>GAL4, UAS-GFP/UAS-Ras<sup>V12</sup>* | 9.5 min<br>10 min | 30 h 25 °C |
| 6 B | *hsflp<sup>122</sup>/+; TRE-RFP/+;*<br>*act>y<sup>+</sup>>GAL4, UAS-GFP/UAS-Ras<sup>V12</sup>* | 9.5 min | 30 h 25 °C |
| 6 C | *hsflp<sup>122</sup>/+; +/+;*<br>*act>y<sup>+</sup>>GAL4, UAS-GFP/+* | 10 min | 30 h 25 °C |
| 6 E | *UAS-fkh-3xHA/hsflp<sup>122</sup>; +/+;*<br>*act>y<sup>+</sup>>GAL4, UAS-GFP/+* | 10 min | 30 h 25 °C |
| 6 F | *UAS-fkh-3xHA/hsflp<sup>122</sup>; +/+;*<br>*act>y<sup>+</sup>>GAL4, UAS-GFP/UAS-Ras<sup>V12</sup>* | 10 min | 30 h 25 °C |
| S6.1 B,F | *hsflp<sup>122</sup>/+; tub-miniCic-mCherry/+; act>y<sup>+</sup>>GAL4,*<br>*UAS-GFP/UAS-Ras<sup>V12</sup>* | 10 min | 30 h 25 °C |
| S6.1 C | *UAS-fkh-3xHA/hsflp<sup>122</sup>; tub-miniCic-mCherry/+;*<br>*act>y<sup>+</sup>>GAL4, UAS-GFP/+* | 10 min | 30 h 25 °C |
| S6.1 D | *UAS-fkh-3xHA/hsflp<sup>122</sup>; tub-miniCic-mCherry/+; act>y<sup>+</sup>>GAL4, UAS-GFP/UAS-Ras<sup>V12</sup>* | 10 min | 30 h 25 °C |
| S6.1 G | *hsflp<sup>122</sup>/+; tub-miniCic-mCherry/UAS-ey; act>y<sup>+</sup>>GAL4, UAS-GFP/ +* | 10 min | 30 h 25 °C |
| S6.1 H | *hsflp<sup>122</sup>/+; tub-miniCic-mCherry/UAS-ey; act>y<sup>+</sup>>GAL4, UAS-GFP/UAS-Ras<sup>V12</sup>* | 10 min | 30 h 25 °C |
| S6.2 A | *hsflp<sup>122</sup>/+; tub-miniCic-mCherry/+; act>y<sup>+</sup>>GAL4, UAS-GFP/UAS-Ras<sup>V12</sup>* | 10 min | 30 h 25 °C |
| S6.2 B | *hsflp<sup>122</sup>/+; tub-miniCic-mCherry/UAS-ey; act>y<sup>+</sup>>GAL4, UAS-GFP/ +* | 10 min | 30 h 25 °C |
| S6.2 C | *hsflp<sup>122</sup>/+; tub-miniCic-mCherry/UAS-ey; act>y<sup>+</sup>>GAL4, UAS-GFP/UAS-Ras<sup>V12</sup>* | 10 min | 30 h 25 °C |

### Immunohistochemistry and imaging

Imaginal discs were dissected and fixed in 4% formaldehyde/PBS for 15 min at room temperature (RT). The samples were washed in 0.1 Triton X-100 (PBT) and then incubated in PBT+5% normal goat serum (PBTN) for 10 min for blocking. Discs were incubated with primary antibodies overnight at 4°C: rabbit anti-Cleaved Drosophila cDcp-1 (1:200, Cell Signaling, #9578S), rabbit anti-fkh (1:200, gift from Martin Juenger), mouse anti-eye (1:100, DSHB anti-eye), mouse anti-wingless (1:100, DSHB #4D4-s), goat anti-Distalless (Santa Cruz Biotechnology Distal-less df-20), rat anti-Ci (1:50, DSHB #2A1-s), rat anti-RFP (1:1000, Chromotek #5F8), rabbit anti-RFP (1:200, MBL #PM005), chicken anti-mCherry (1:1000, Abcam #ab205402), rat anti-Drosophila E-cadherin (1:50, DSHB DCAD2-s), chicken anti-GFP (1:1000, Abcam #ab13970). The discs were then washed in PBT, followed by another blocking step with PBTN. The samples were counterstained with: DAPI (0.25 ng/l, Sigma), Phalloidin (Abcam Phalloidin 405 ab176752 1:1000, Sigma Aldrich Phalloidin 555 P1951 1:400, Invitrogen Phalloidin 647 1:100) and secondary antibodies from Invitrogen, 1:500 (goat anti-rabbit A11008, goat anti-chicken A11039, goat anti-chicken A21437, goat anti-rat A21434, goat anti-mouse A32728, goat anti-rabbit A21244, goat anti-chicken A21449, goat anti-mouse A21235, donkey anti-goat A32849) and incubated for 3 h at room temperature. Discs were again washed in PBT and PBS, then mounted using Molecular Probes Antifade Reagents (#S2828). To prevent squeezing of samples by coverslips for imaging the apical architecture without interference from the peripodium (Fig. S5.1), two stripes of double-sided tape (Tesa, #05338) were placed on the slide. Samples were imaged using a Leica SP8 confocal microscope. The figures were assembled in Affinity Design.

### Image analysis and statistics

Where possible, control and experimental samples were fixed, processed and mounted together to ensure comparable staining and imaging conditions. The signals of the following fluorescent reporters were further amplified by anti-GFP or anti-RFP antibody staining: *TRE-RFP* and *miniCiC-mCherry*. Images were processed, analysed and quantified using tools in Fiji (ImageJ2.3.0/) [79] (see details below). Great care was taken to apply consistent methods (i.e. number of projected sections or thresholding methods) within experiments. Statistical tests were performed in GraphPad PRISM9. Details and the number of wing discs used for each test (n), can be found in the respective figure legends. Figure panels were assembled using Affinity Design.

### Image segmentation and quantification

#### Epithelial integration of clones

Z-projections of maximum intensity of 2-3 slices of apical and basal sections were generated. A ROI was defined around the central region of the pouch, to avoid confounding results due to the folded structure of the wing disc in the hinge. The number of clones in the ROI in the pouch of the apical and basal sections were counted. A paired t-test was used to statistically compare the number of clones detected apically and basally.

#### Quantification of apoptosis within clones

Z-projections of maximum intensity of 2-3 slices of basal sections were generated. To define the total region of the wing disc, a Gaussian blur filter of sigma=4 was applied to the DAPI channel. Intensity-based thresholding using ‘triangle dark’ threshold function was then used to generate a DAPI-based wing disc mask. To create a mask of GFP-labelled clones, the variation in GFP intensities were first pre-processed by applying a maximum and minimum filter of radius=2. Then, intensity-based thresholding using the ‘default dark’ threshold function was performed to create a binary image. ‘despeckle’ and ‘fill holes’ functions as well as a size inclusion range of 10-infinity µm^2^ was applied to create the final clonal mask. The wing disc mask and the clonal GFP mask were used to define (by Boolean functions) different ROIs and then extract parameters, such as total clonal area, whole disc area, and background area. To define the apoptotic area, the cDcp1 channel was pre-processed by applying maximum and minimum filters, each of radius=1. An intensity threshold (‘intermodes dark’) for cDcp1 channel was set in such a way that only high intensity cDcp1 particles were picked up. cDcp1 particles were quantified in the different ROIs by ‘limit to threshold function’, and the percentage apoptotic area in each ROI was calculated. A one-way ANOVA test was performed to test for statistical significance.

#### Quantification of TRE-RFP reporter activity in interface contractility and cell-cell competition models

Z-projections of maximum intensity of 2-3 slices of basal sections were generated. To define the total region of the wing disc, a Gaussian blur filter of radius=4 was applied to the DAPI channel. Intensity-based thresholding using ‘minimum dark’ threshold function was then used to generate a DAPI-based wing disc mask. To create a mask of GFP-labelled clones, the variation in GFP intensities were first pre-processed by applying a maximum and minimum filter of radius=2. Then, intensity-based thresholding using the ‘otsu’ threshold function was performed to create a binary image. ‘despeckle’ and ‘fill holes’ function as well as a size inclusion range of 10-infinity µm^2^ was applied to create the final clonal mask. To determine the average size of cells in the wing disc to define a 1-cell wide ROIs for wild type and clonal interfaces, we measured the size of 30 nuclei in 3 wild type wing discs, and determined their average size as 3.75µm. To account for cytoplasm in the densely packed pseudostratified tissue, we set the size of each ROI to be 4µm wide. The enlarge function was used on the GFP clonal mask to generate a 4µm thick band around clones, representing the wild-type interface ROI (dark blue) as an approximately 1-cell thick band. Then, a second mask of only large clones was generated by defining a ROI with clones of a minimum area of 180 µm^2^ to visualize and only select clones that also contained an ‘interior clonal cells (light pink)’ population. On this second mask of large clones, we applied the enlarge function of −4 µm to create ‘clonal interface cells’ ROI (magenta) and the ‘interior clonal cells’ ROI (light pink). Using Boolean functions on these ROIs, the wing disc and the clonal mask, all ROIs required for further analysis could be generated. For example, to generate the background wild type cell ROI, clones and their wild type interface regions were excluded from the whole disc. Ultimately, the fluorescence intensity of the TRE-RFP channel was measured in different ROIs. Please note that for the quantification of the background TRE-RFP intensity in the discs with RpS3^-/-^ clones (Fig 4), the background was defined as a 4µm band of wild type cells adjacent to the wild type interface cells, as we observed a general upregulation of JNK-signalling independent from an interface pattern in the mosaic RpS3^-/-^ wing discs. One-way ANOVA tests were performed.

#### Quantification of TRE-RFP reporter activity in the centre and periphery of the pouch in clones expressing GFP or UAS-Tkv^CA^ clones

A single representative section of the disc proper was selected, which contained nuclei of both clonal cells and wild type cells and used for further analysis. To create a mask of GFP-labelled clones, the variation in GFP intensities were first pre-processed by applying a maximum and minimum filter of radius=2. Then, intensity-based thresholding using the ‘otsu’ threshold function was performed to create a binary image and a size inclusion range of 10-infinity µm^2^ was applied to create the final clonal mask. Individual clones were the picked in such a way that a 4 µm band could be drawn around them using the enlarge function, without this region encroaching into the area of surrounding clones. The criteria for picking different clones were as follows: *Dad*-LacZ intensity in clones in the centre of the pouch and in the 4 micron band around the respective clone was in the range of +-20AU; however, the difference in intensity in *Dad*-lacZ clones and the 4-micron band around the clone was much larger in the pouch periphery (the Dad-LacZ intensity in the 4-micron band around clones in pouch periphery was close to 0). The channel with TRE-RFP signal was selected and the fluorescence intensity in the clones in the centre and periphery of the pouch were measured. A paired t-test was used to statistically analyse the TRE-RFP reporter intensity in clones expressing GFP or Tkv^CA^ in the centre and the periphery of the pouch.

#### Quantification of percentage of apoptotic area in the presence and absence of JNK signalling

The percentage of apoptotic area in wing discs expressing aberrant clones was analysed in in presence (5B, 5C S5B, S5C) and in the absence of JNK signalling (5H,L). We used two different model systems to study the effect of JNK inhibition on the apoptosis of aberrant clones: We first suppressed intra-clonal JNK activity only, for example flip-out clones co-expressing UAS-tkv^CA^ and UAS-bsk^DN^ (5H). For the second model, we suppressed JNK activity in the entire posterior compartment using the en-GAL4-UAS expression system, and the tkv^CA^ expressing clones were under the control of the LexA-LexO system (5L).

Z-projections of maximum intensity of 5-6 slices of basal sections were generated. To define the total region of the wing disc, a Gaussian blur filter of radius=4 was applied to the DAPI channel. Intensity-based thresholding using ‘minimum dark’ threshold function was then used to generate a DAPI-based wing disc mask. To create a mask of GFP-labelled clones, the variation in GFP intensities were first pre-processed by applying a maximum and minimum filter of radius=2. Then, intensity-based thresholding using the ‘otsu’ threshold function was performed to create a binary image. ‘despeckle’ and ‘fill holes’ functions as well as a size inclusion range of 10-infinity µm^2^ was applied to create the final clonal mask. The enlarge function was used on the GFP clonal mask to generate a 4µm thick band around clones, representing the wild type interface ROI (dark blue) as an approximately 1-cell thick band around every clone. Then, a second mask of only large clones was generated by defining a ROI with clones of a minimum area of 180 µm^2^ in to visualize and only select clones that also contained an ‘interior clonal cells (light pink)’ population. On this second mask of large clones, we applied the enlarge function of −4 µm to create ‘clonal interface cells’ ROI (magenta) and the ‘interior clonal cells’ ROI (light pink). Additionally in 5H, ROIs of the posterior and anterior wing disc compartments were generated using the en-GAL4,UAS-GFP channel and the Boolean function XOR, respectively. Using Boolean functions on these ROIs, the wing disc and the clonal mask, all ROIs required for further analysis could be generated. For example, to generate the background wild type cell ROI, clones and their wild type interface regions were excluded from the whole disc. To define the apoptotic area, the cDcp1 channel was pre-processed by applying maximum and minimum filters, each of radius=1. An intensity threshold (‘intermodes dark’) for cDcp1 channel was set in such a way that only high intensity cDcp1 particles were picked up. cDcp1 particles were quantified in the different ROIs by ‘limit to threshold function’, and the percentage apoptotic area in each ROI was calculated. A one-way ANOVA test was performed to test for statistical significance.

#### Analysis of spatial correlation between buckling and apoptosis

Maximum intensity projections were generated, which specifically excluded the peripodium, when analysis of all apoptotic events in 3D was required. The GFP-marked clonal mask was generated using intensity-based thresholding, followed the ‘fill holes’ and ‘despeckle’ functions to remove noise. Medium-sized clones with an area of buckling as well as a planar area were selected and added to the ROI manager. A whole disc cDcp1 mask was generated using intensity-based thresholding on the cDcp1 channel. A clonal cDcp1 mask was generated using Boolean function AND, and added to the ROI manager. Maximum projections of the apical junctional network (E-cad) were used to outline the buckling area within a clone with the oval selection tool, which was then added to the ROI manager. Buckling was defined as a characteristic reduction in cell surface areas observed in max projections of the curved surface during buckling. Buckling was verified by analysis of the reprojected Z-sliced stack. An area of similar size and shape was then selected in non-buckling, planar area of the same clone. Care was taken that the planar region was not close to a clonal boundary, to avoid confounding effects of JNK-dependent interface signalling on apoptotic behaviour. The clonal cDcp1 within the buckling area or the planar area were combined by means of the ‘AND’ function. Lastly, cDcp1 area measurements in both buckling and planar areas were performed. 11 clones from at least three different wing discs were quantified.

#### Quantification of *miniCic*-reporter activity

A single representative section of the disc proper was selected from the stack for further analysis. A nuclear binary mask for the whole disc was obtained by intensity-based thresholding of the DAPI channel and added to the ROI manager. To create a binary mask of GFP-labelled clones, we used intensity-based thresholding, followed by the ‘despeckle’ function and ‘fill holes’ function. To compare the clonal *miniCic* signal with the signal in surrounding WT cells, individual clones were selected as and added to the ROI manager. An outer band ROI (4 μm, corresponding to one cell row) was established around the selected clones using the ‘enlarge’ function. A ROI of clonal nuclei was obtained by using the Boolean function AND on the nuclear mask and the respective clone ROI. Similarly, the outer band of nuclei ROI was also generated for the respective clones. Then the fluorescence signal intensity in the *miniCic* channel was measured for individual clones in the clonal nuclei ROI and the outer band nuclei ROI. The number of clones analysed in each experiment is noted in the figure legends.

---

**Figure S1.**
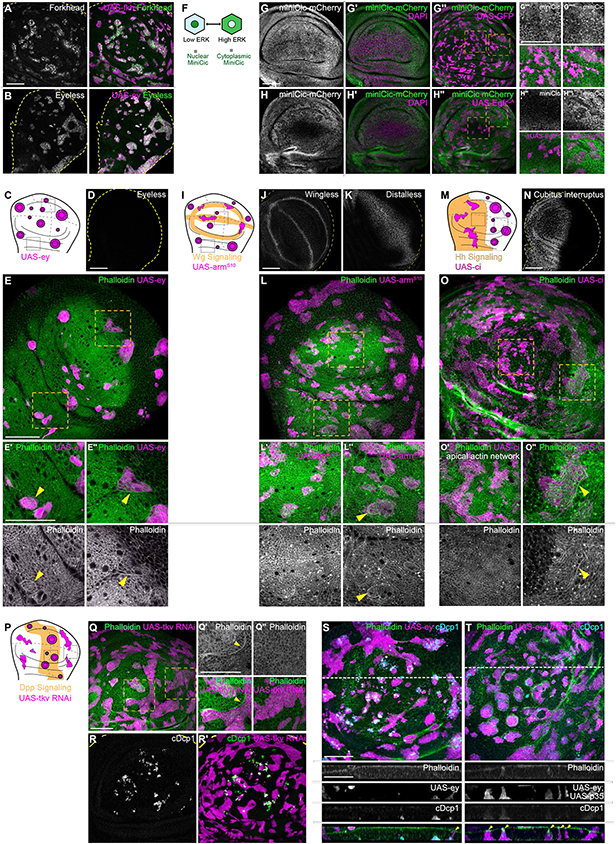
Apoptosis is essential to eliminate cells by interface contractility. **A,B**: Wing discs carrying mosaic clones (magenta) expressing Fkh (A) or Eyeless (B), were stained for Fkh (A) and Ey (B) proteins, respectively, to validate anti-Fkh and anti-Ey antibodies. Dashed yellow lines demarcate wing disc boundaries. **C, I, M:** : Wing discs schemes illustrating endogenous patterns of Ey expression (C), Wg signalling (I) or Hh signalling (M) (orange). Clones that do not induce interface contractility, because they only express GFP or whose fate is like that of surrounding cells (magenta clones in orange domains in I, M), maintain irregular clone shapes. Clones whose fate is very different to surrounding cells, because of changes to cell fate and differentiation programs (magenta clones in white domains in C, I, M) experience interface smoothening and even cyst formation. Clones express Ey (C), Arm^S10^ (I), or Ci (M), activating Ey, Wg or Hh cell fate programs, respectively. **F**: Scheme to illustrate miniCic reporter activity. Nuclear localization of the miniCic reporter indicates low ERK signalling and cytoplasmic localization indicates high ERK signalling. **G,H**: Wing disc expressing the miniCic-mCherry reporter (grey or green, G-G’’’’,H-H’’’’) ubiquitously and carrying mosaic clones expressing GFP (magenta, G’’-G’’’’) or Egfr^CA^ (magenta, H’’-H’’’’) in the centre (’’’) and periphery (’’’’) of the pouch. Discs were stained with DAPI to visualize nuclei (grey or magenta in ‘). Yellow frames mark regions shown in (’’’and’’’’) panels. **D, J, K, N** : Wing discs were stained to visualize expression patterns of the transcription factor Ey (D), the morphogen Wingless (J), the Wg target gene Distalless (K) and the Hh-effector Ci (N). Dotted yellow lines demarcate wing disc boundaries. **E, L, O**: Wing disc carrying mosaic clones (magenta) that express Ey (E), Arm^S10^ (L) or Ci (O) were stained with phalloidin to visualize Actin (grey or green). Yellow frames mark regions in pouch centre (E’, L’), pouch periphery (E’’), hinge (L’’), anterior compartment (O’), and posterior compartment (O’’). Please note: The images in O’ are from a section more apical to the one shown in O. This was done in order to properly view the apical phalloidin network in the anterior compartment. **P:** Wing disc schemes illustrating endogenous patterns of Dpp signalling (orange). Clones that express Tkv RNAi do not induce interface contractility in peripheral domains (white), where Dpp signalling is low. The fate of tkv RNAi expressing clones is different to surrounding cells in high Dpp signalling domains (orange). Therefore clones experience interface smoothening and even cyst formation. **Q:** Maximum intensity projection of apical sections of a wing disc carrying mosaic clones (magenta) expressing a Tkv RNAi construct, stained with phalloidin to visualize Actin (grey or green). Clone smoothening and actin enrichment (yellow arrow) in areas of Dpp signalling (Q’). Irregular clone shapes are maintained in regions without Dpp signalling (Q’’). Yellow frames mark regions shown in (Q‘ and Q’’) panels. **R**: Maximum intensity projection of basal sections of a wing disc carrying mosaic clones (magenta) expressing a Tkv RNAi construct, stained for cDcp1 to visualize apoptosis (grey or green). Dashed yellow lines demarcate wing disc boundaries. **S, T**: Maximum-intensity projections of basal sections of wing discs carrying clones (magenta) expressing Ey (S), or Ey;p35 (T), were stained with phalloidin to visualize Actin (grey or green) and for cDcp1 to visualize apoptosis (grey or cyan). Dashed white lines indicate position of cross-sections. Scale bar = 50µm.

**Figure S2.1.**
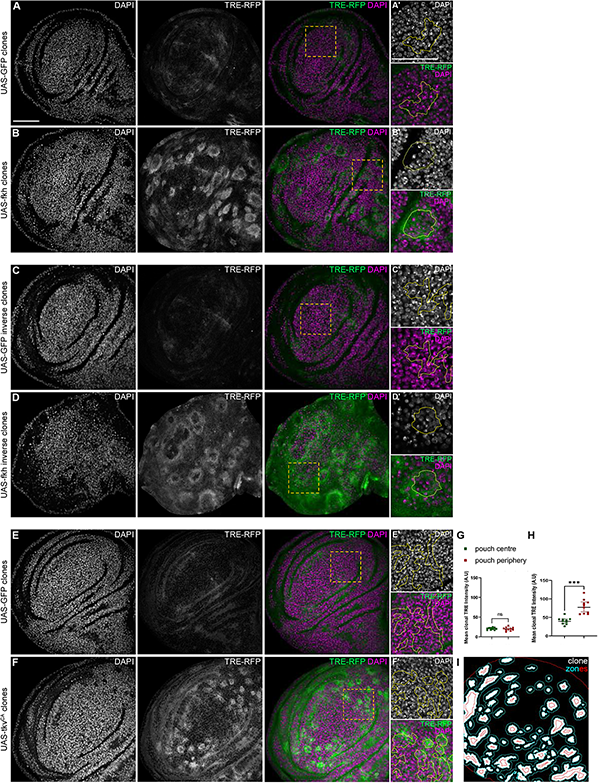
JNK activation occurs in both sides of interface-contacting cells. DAPI images form main figure panels are shown to validate the conclusion that signalling at the interface affects a single row of cells on each side. For clonal marker visualization, please refer to the main Fig.2. **A-F**: Lateral sections of wing imaginal discs expressing the TRE-RFP reporter (grey or green) and carrying mosaic clones (grey in Fig.2 A, D, F, H, K, L) expressing GFP (A, C, E) and Fkh (B,D) or Tkv^CA^ (F). Discs were stained with DAPI to visualize nuclei (grey or magenta). Yellow frames mark regions shown in in (‘and’’) panels. Continuous yellow lines in (‘and’’) panels mark clone boundaries. **G,H**: Quantifications of TRE-RFP reporter intensity in clones expressing GFP (G) or Tkv^CA^ (H) when located either in the central domain of the pouch where Dad-LacZ expression is normally high, or in the pouch periphery, where Dad-LacZ expression is normally low. Graphs display mean± 95% CI. Statistical analysis was done using Paired Student’s T-tests with ns = not significant, *** p≤0.001. n=10 clones in the centre of the pouch, and n=10 clones in the periphery of the pouch, per genotype. **I**: Example of a segmentation mask based on clone areas (white), clone interfaces and adjacent 4µm zones, which were used to quantify TRE-RFP reporter intensities. Scale bar = 50µm

**Figure S2.2.**
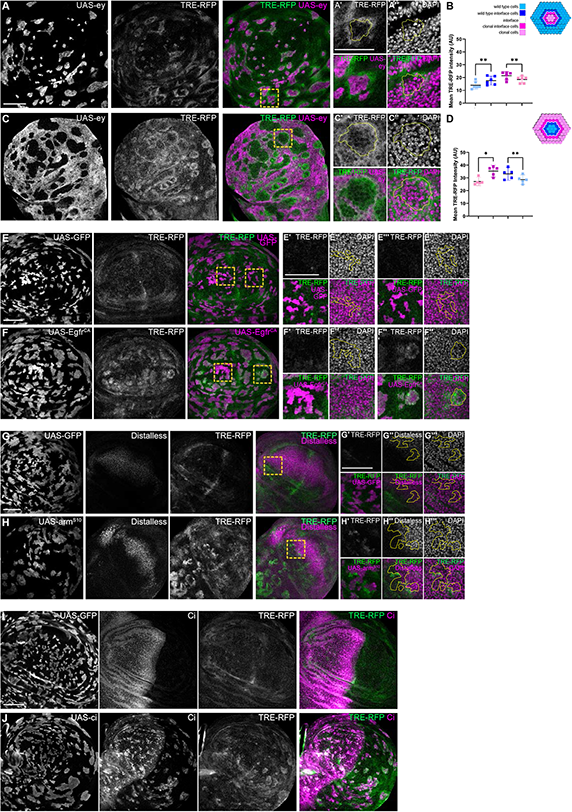
JNK interface signalling is a robust hallmark of interface contractility responses. **A,C**: Lateral section of a wing disc expressing the TRE-RFP reporter (grey or green) and carrying mosaic clones expressing Ey (grey or magenta) as minority (A) or majority (C) in the wing disc. Different length of heat shock induction allows scaling of clone sizes. Yellow frames mark regions shown in (A’, A’’, C’, C’’). Continuous yellow lines in (‘ and “) panels represent clone boundaries. **B,D**: Quantifications of TRE-RFP intensities in specific zones of clones expressing Ey, if they represent the minority (B) or majority (D) of cells in the disc. One-way ANOVA tests were performed to test for statistical significance, ns = not significant, *0.05>p, **0.01>p. n=6 wing discs (B) and n=5 wing discs (D). Schemes depicting specific zones in and around clones that were quantified are provided next to the respective graphs. **E,F:** Lateral section of a wing disc expressing the TRE-RFP reporter (grey or green) and carrying mosaic clones (magenta) expressing GFP (E) or Egfr^CA^ (F). Yellow frames mark regions shown in (‘-’’’’) panels. Yellow lines in (‘-’’’’) panels demarcate clone boundaries. **G,H**: Lateral section of a wing disc expressing the TRE-RFP reporter (grey or green) and carrying mosaic clones (magenta) expressing GFP (G), and Arm^S10^ (H). Discs were stained for Distalless, to visualize one target gene of Wg-signalling. Yellow frames mark regions shown in G’, G’’, G’’’, H’, H’’, H’’’. Yellow lines in (‘-’’’) panels demarcate clone boundaries. **I,J**: Lateral section of a wing disc expressing the TRE-RFP reporter (grey or green) and carrying mosaic clones (magenta) expressing GFP (I), or Ci (J). Discs were stained for Ci to visualize relative Ci expression and thus patterning differences. Scale bar = 50µm

**Figure S2.3.**
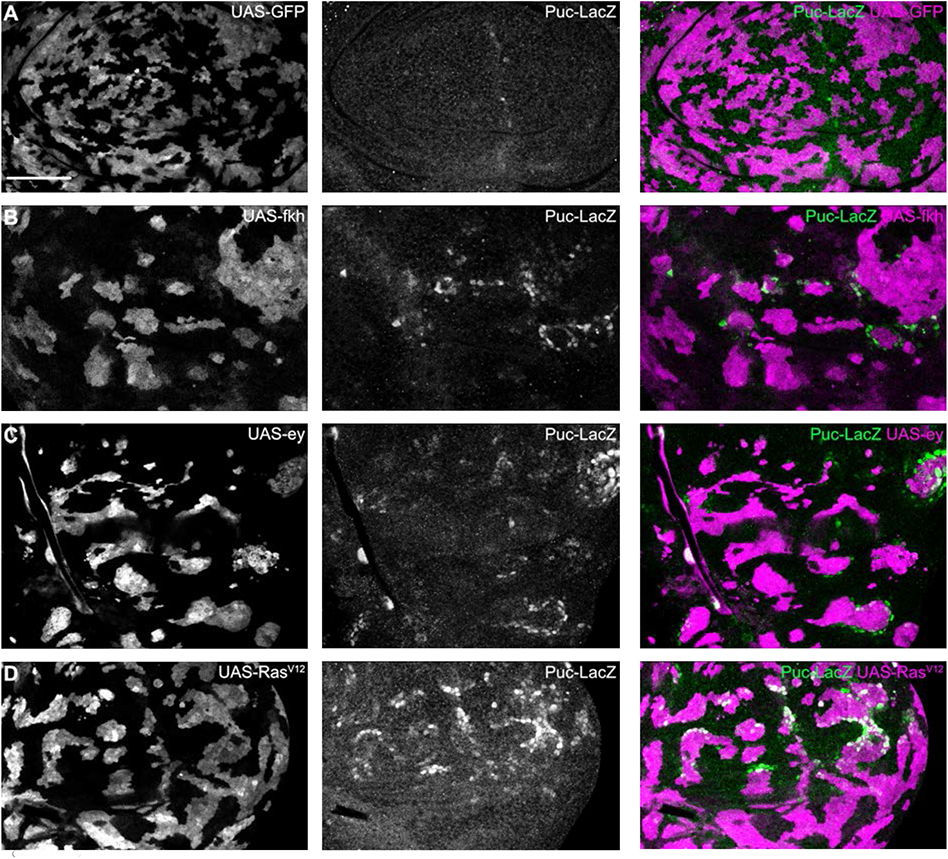
Interface JNK signalling can be detected using the puc-LacZ reporter. **A-D**: Lateral section of wing discs expressing the puc-LacZ JNK reporter (grey or green), and carrying mosaic clones (grey or magenta) expressing GFP (A), Fkh (B), Ey (C), or Ras^V12^ (D). Scale bar = 50µm.

**Figure S3.**
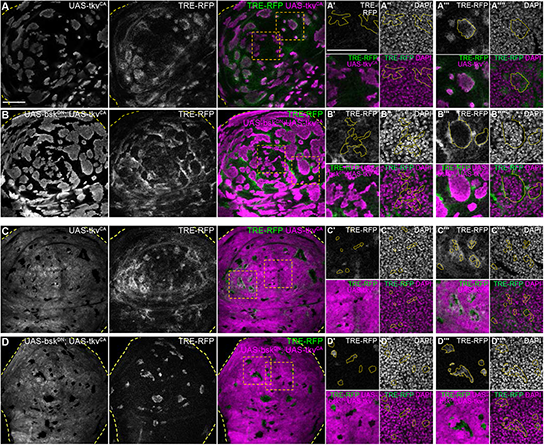
JNK is activated cell-autonomously at clonal interfaces. **A-D:** Lateral sections of wing discs expressing the TRE-RFP reporter (grey or green) and carrying mosaic clones (grey or magenta) expressing Tkv^CA^ (A,C), or Bsk^DN^, Tkv^CA^ (B,D). Discs were stained with DAPI to visualize nuclei (grey or magenta) in (“ and “”) panels. Dashed yellow lines demarcate wing disc boundaries. Yellow frames mark regions shown in (‘-’’’’) panels. Yellow lines in (‘-’’’’) panels demarcate clone boundaries. Scale bar = 50µm

**Figure S4.1.**
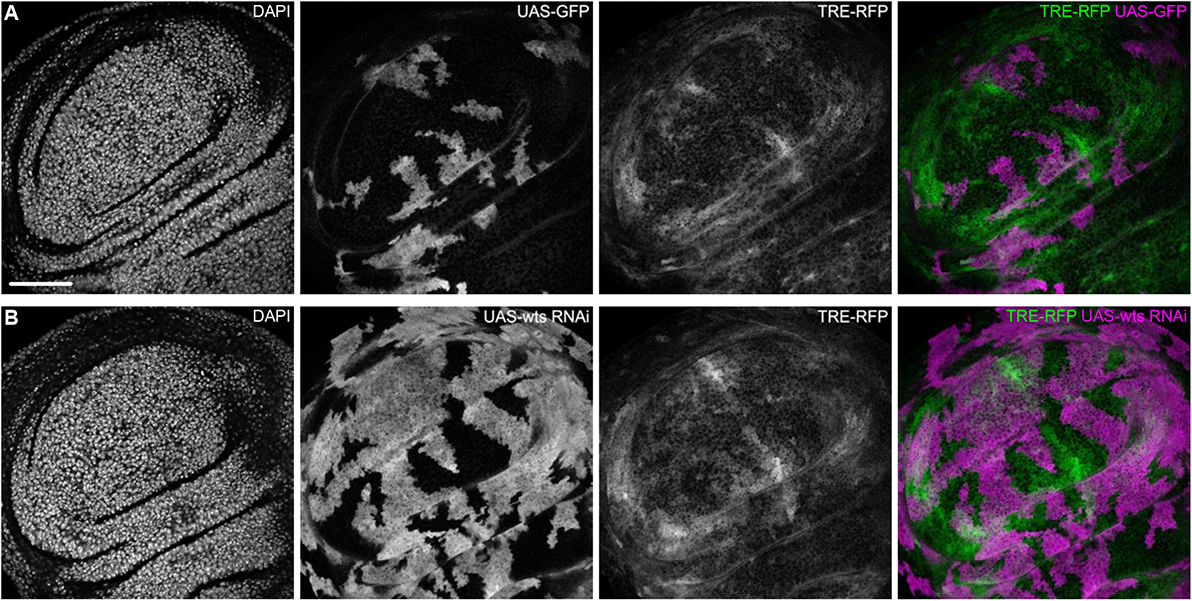
JNK interface signalling is unique to interface contractility. **A,B:** Lateral sections of wing discs expressing the TRE-RFP reporter (grey or green) and carrying mosaic clones (grey or magenta) expressing GFP (A), or a *wts* RNAi construct (B). Scale bar = 50µm

**Figure S4.2.**
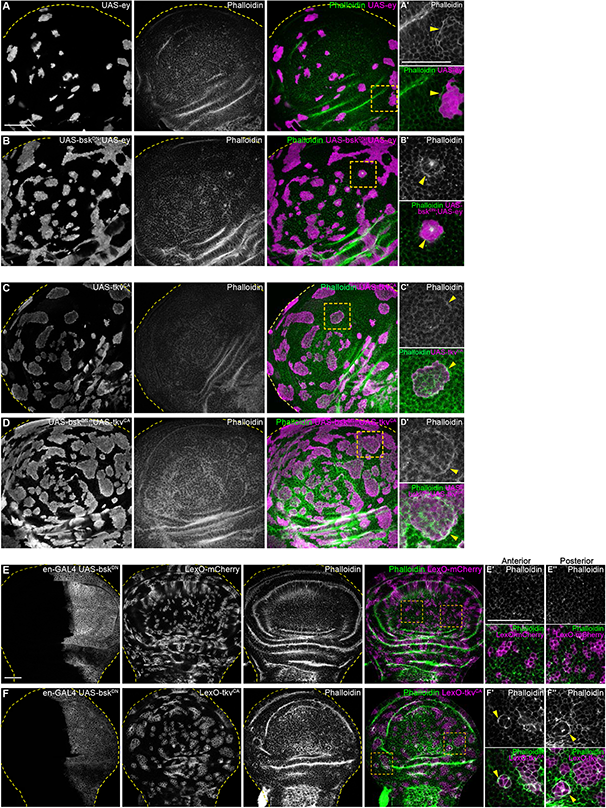
JNK signalling is not required for actomyosin enrichment at clonal interfaces. **A,B,C,D:** Lateral sections of wing discs carrying mosaic clones (grey or magenta) expressing Ey (A), Tkv^CA^ (C) or Bsk^DN^, Ey (B), Bsk^DN^, Tkv^CA^ (D). Discs were stained with Phalloidin to visualize Actin (grey or green). Dashed yellow lines demarcate wing disc boundaries. Yellow frames mark regions shown in A’, B’. Yellow arrow heads highlight actin enrichment at the clone boundary (A’, B’). **E, F:** Lateral sections of wing discs where the posterior compartment expresses Bsk^DN^ under the control of *en*-GAL4 (grey E,F), and carrying mosaic clones (grey or magenta) expressing LexO-mCherry (E’, E’’) and LexO-Tkv^CA^ (F’, F’’). Discs were stained with phalloidin to visualize Actin (grey or green). Dashed yellow lines demarcate wing disc boundaries. Yellow frames mark regions shown in the anterior (E’,F’) and posterior (E’’, F’’) compartment. Yellow arrow heads in (‘ and “) panels point to actin enrichment at the clone boundary. Scale bar = 50µm

**Figure S5.**
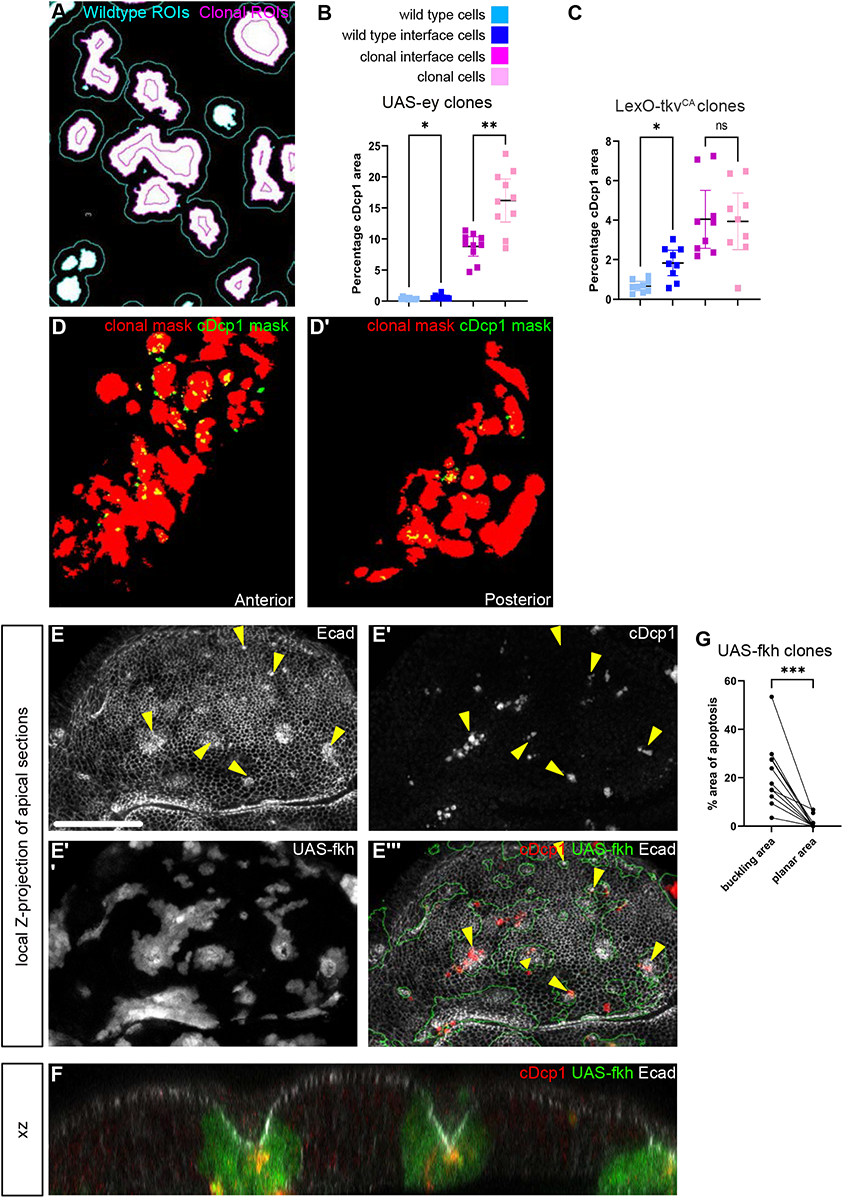
Apoptosis can be observed at the buckling points of Fkh expressing clones. **A:** Example of a segmentation mask based on clone areas (white), and ROIs corresponding to wildtype interface cells (cyan) and clonal interface cells (magenta). **B,C**: Quantifications of cDcp1 area fractions in selected zones around clones expressing UAS-Fkh (B) or UAS-Tkv^CA^ (C). Only large clones with distinct interior zones were quantified for clone interior and clonal interface zones. Graphs display mean± 95% CI. One-way ANOVA tests were performed to test for statistical significance, ns = not significant, * p≤0.05, ** p≤0.01, *** p≤0.001. n=12 wing discs (B) and n=8 wing discs (C). **D:** Overlay of segmentation mask of LexO-Tkv^CA^ clones (red) and cDcp1 (green) in anterior (B) and posterior (B’) compartments, in a wing disc where the entire posterior compartment expressed UAS-Bsk^DN^. **E:** Local z-projection of apical sections of a wing disc carrying mosaic clones (grey or green outline) expressing Fkh. Discs were stained for cDcp1 to visualize apoptosis (grey or red) and for E-cadherin to visualize the apical junctional network (grey). Yellow arrows point to positions of apical deformations arising from buckling and cystic invagination. **F:** XZ cross-section through a wing disc carrying mosaic clones expressing Fkh (green). The disc was stained for cDcp1 to visualize apoptosis (red) and for E-cadherin to visualize the apical junctional network (grey). **G:** Quantification of the percentage of cDcp1 area found in domains with buckling points versus planar regions within Fkh-expressing clones. A paired Student‘s t-Test was performed to test for statistical significance, *** p≤0.001. n=11 buckling and n=11 planar regions, each pair within the same clone, in 3 wing discs. Scale bar = 50µm

**Figure S6.1.**
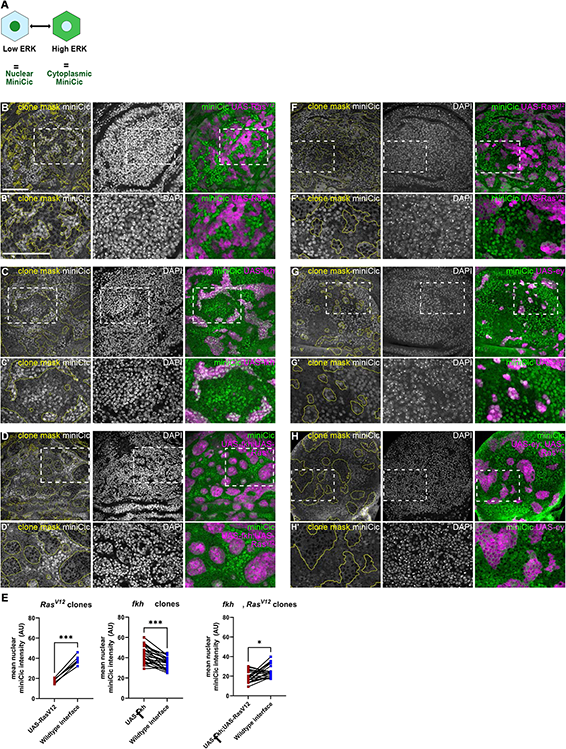
Ras^V12^ dominantly induces high ERK signalling in Fkh- and Ey-expressing clones. **A**: Scheme illustrating *miniCic* reporter activity. Nuclear localization of the *miniCic* reporter indicates low ERK signalling and cytoplasmic localization indicates high ERK signalling. **B-D, F-H:** Lateral section of wing discs expressing the ERK reporter *miniCic*-mScarlett or *miniCic*-mCherry (grey or green) and carrying mosaic clones (yellow outline or magenta) expressing Ras^V12^ (A, F), Fkh (B), Fkh,Ras^V12^ (C), Ey (G) and Ey,Ras^V12^ (H). Discs were stained with DAPI to visualize nuclei (grey). White frames mark regions shown in B’-D’ and F’-H’. **E**: Quantifications of nuclear *miniCic* intensity (AU) in clones expressing Ras^V12^, Fkh or Fkh, Ras^V12^ (dark red) versus nuclear *miniCic* intensity (AU) in corresponding wildtype interface cells (blue). Paired Student’s T-tests were performed to test for statistical significance, * p≤0.05, *** p≤0.001. Scale bar=50µm.

**Figure S6.2.**
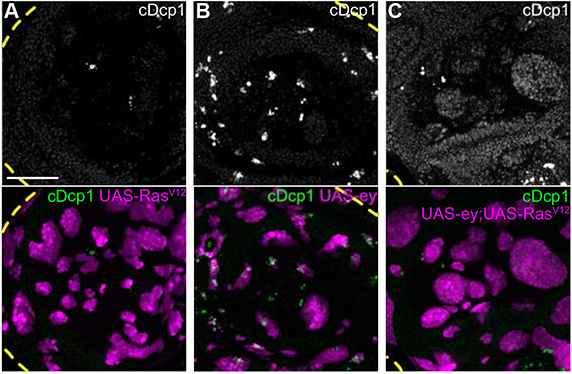
Ras^V12^ rescues Ey-expressing clones from apoptosis. **A,B,C:** Maximum intensity projection of basal sections of wing discs carrying mosaic clones (magenta) expressing Ras^V12^ (A), or Ey (B), or Ey,Ras^V12^ (C). Discs were stained for cDcp1 (grey or green) to visualize apoptosis.). Dashed yellow lines demarcate wing disc boundaries. Scale bar = 50µm

